# Parallel Networks to simulate complex multicellular dynamics - A proof of concept with intervertebral disc cell systems

**DOI:** 10.1101/2022.08.08.503195

**Authors:** Laura Baumgartner, Miguel Ángel González Ballester, Jérôme Noailly

## Abstract

**Background:** Network models are convenient to represent in a mechanistic way the complexity of cell biological activity. Dynamic simulations of such networks might require approximations of equation parameters through reverse engineering, numerous and costly experimental research, and/or have limited capacity to explore cell responses to chronic, dose-dependent stimulus exposure. Here we present a mechanistic methodology allowing the simulation of interrelated cell responses of multicellular systems to multifactorial stimuli with dose-and time dependent network links.

**Methods:** a mathematical framework to approach systems biology research questions is presented, the Parallel Networks (PN)-Methodology. It consists of a novel concept, where multicellular systems are described as many relatively small feed-forward networks acting in parallel. Each parallel network is calculated through a specifically designed ordinary differential equation (ODE). Through a unique approach to feed the ODE with interrelated parameters, a system of decoupled, analytically resolvable ODE was obtained.

**Results:** Applied to intervertebral disc multicellular systems, virtual environments of multifactorial stimuli and multiple cell responses simulating daily moving activities, and to microgravity could successfully be created.

**Conclusions:** The PN-Methodology stands for a one-of-a-kind mathematical methodology to approximate dynamics of complex multicellular systems over long periods of time at low computational costs.

## 1. Introduction

Advances in experimental and *in silico* research importantly enhance our understanding of biological systems. In *in silico* research dynamics of biological phenomena are approximated through mathematical concepts. An important research area is graph-based (network) modeling, where targets (e.g. organs, genes or proteins) are represented as nodes that are linked among each other(1). In the case of oriented networks, the links usually represent the activating or inhibiting relationships among targets, providing a certain degree of mechanistic understanding of the model (1,2). Network models can be built for any spatial scale with either a bottom-up or a top-down approach. At the (sub-) cellular level, bottom-up descriptions are more common. Thereby, (intra-)cellular components and their interactions are simulated to approximate cell responses. This classically includes protein-protein interaction networks, cell signaling networks, metabolic networks or gene regulatory networks (3). Known interactions are often available through curated databases. An overview of publicly available databases on e.g. protein-protein interactions and signaling pathways was previously provided (4). In general there is a need for finding proper guesses in terms of network topology and directionality. The corresponding exploration of the best network configurations might come from numerous, cost and time intensive (biochemical) experiments or are obtained by reverse engineering. The latter is usually coupled to demanding optimization problems within a high-dimensional parameter space, where heuristic methods and/ or, iterative algorithms shall be required to find satisfying solutions (5).

Mathematical solutions to determine network dynamics can roughly be divided into non-deterministic (probabilistic) and deterministic, even though methods that combine these approaches also exist (6). Options to obtain a deterministic response of a network include ordinary differential equation (ODE)-based approaches and logical modeling. ODE-based solutions usually require resolving systems of ODE. ODE solvers, however, need to define crucial mathematical criteria for the calculation of the relative and absolute errors and tolerances, which affects network results (7). Logical modeling only requires information about the qualitative nature (usually activating or inhibiting) and the directionality of links within a network, hence, requiring less information than quantitative ODE-based approaches. Accordingly, network responses are rather qualitative (Boolean) or semi-quantitative (fuzzy logic), usually leading to attractors represented by steady state nodal activations under constant initial network perturbations. Hence, logical models often lack sensitivity to time and to the dose of a node, which limits the estimates of the evolution of dynamical systems over time, where the qualitative nature of a link (i.e. activating or inhibiting) might change.

The simulation of the changing nature of a stimulus due to the stimulus dose and along time is cornerstone, however, in the exploration of the long-term turnover of human body tissues, in health and disease, and holds particularly true for the simulation of load-bearing organs (e.g. (8,9)). Considering that tissue dysfunctions might develop slowly, i.e., over months, years and decades, the computational efficiency of time-dependent mathematical solutions, further stands for a key technical requirement.

To investigate dysfunctions of mechanosensitive tissues, relative responses of cells within multicellular systems are of interest, which are exposed to highly complex, multifactorial stimulus environments.

Accordingly, we propose a novel concept in network modeling: the “parallel networks (PN)”. This is introduced as a high-level concept to comprehensively estimate dynamics of multicellular environments whilst allowing for changes of the nature of the links according to the dose and exposure time of the cell stimuli. This novel mathematical framework is presented as the “PN-Methodology”. Its position among currently existing numerical modeling approaches in engineering and biology is illustrated in Figure 1.

**Figure 1:**
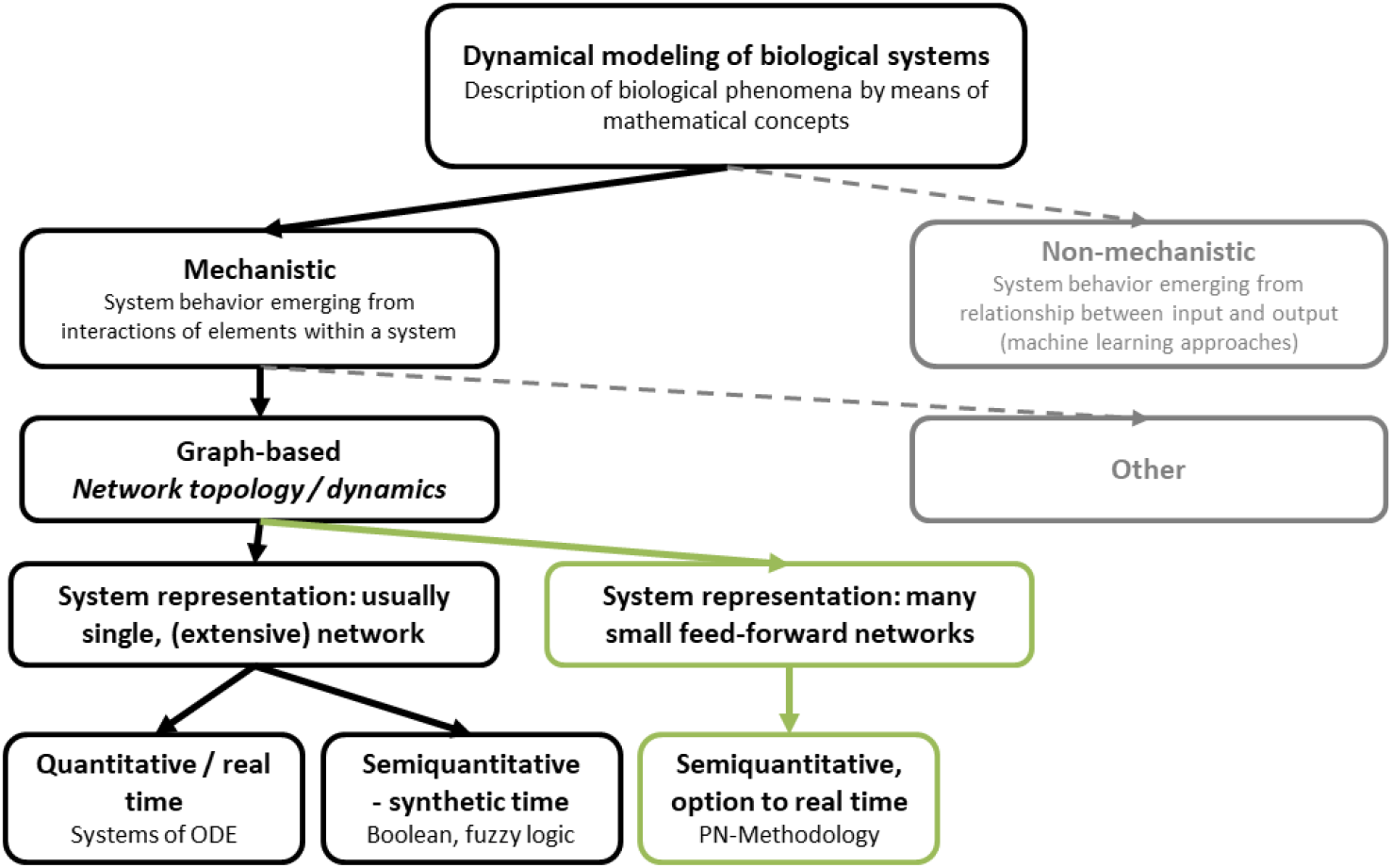
The PN-Methodology (green) situated within the mechanistic *in silico* (computer) modeling biology and engineering domain.

Intervertebral disc (IVD) degenerative changes were simulated as an illustrative case of multifactorial disorder that affects a load-bearing organ over long periods of time. The IVD provides flexibility to the spine and contributes to the mechanical stability of the trunk. At the same time, it is the biggest avascular structure of the human body. Therefore, disc cells, especially the ones of the Nucleus Pulposus (NP), i.e., the center of the IVD, are continuously exposed to changing mechanical loads, and to critical nutritional conditions. The latter is, amongst others, characterized by low glucose (glc) concentrations and by an acidic environment (4). Experimental studies showed that both, adverse nutritional environments and adverse loading conditions can provoke catabolic shifts in disc cell activity (CA) (10–15), and that CA depends on the exposure time to loading conditions (16). By simulating NP cell dynamics in different combinations of metabolic, inflammatory and mechanical environments along time, we could demonstrate that the PN-Methodology provides excellent qualitative forward estimates of the CA within the multicellular NP system.

## 2. Methods

### 2.1 Overview of the PN-Methodology as a high-level concept

The PN-Methodology interprets each cell response to a current stimulus environment as one parallel network, or PN. Hence, many cell responses are considered as many networks acting in parallel. Eventually, many relatively small and feedforward networks are interrelated to describe a complex biological system, and they are dynamically calculated with a specifically designed non-autonomous, linear ODE; the PN-Equation. To build the PN topology and feed the PN-Equations, data- or knowledge-based evidence are used, about cell responses to variations of key relevant stimuli. This allows to treat the cell as a “black box”, as far as intracellular regulation is concerned. In other words, “final” cell biology data of interest such as cell transcriptomics or proteomics are targeted and semi-quantitatively calculated according to environmental stimulus concentrations.

The hypothesis behind this approach his that alterations in CA are not spontaneous processes, but initially caused by alterations of external stimuli, e.g. nutrition-related or mechanical stimuli. External stimuli subsequently regulate the CA, both directly, and indirectly, through local stimulus expressions that further alter the CA within the region.

Thus, the methodology allows to consider different cell states (CS) caused by locally different cell environments within a multicellular system, which we call “heterogeneous” stimulus environment. The workflow of the PN-Methodology is shown in Figure 2.

**Figure 2:**
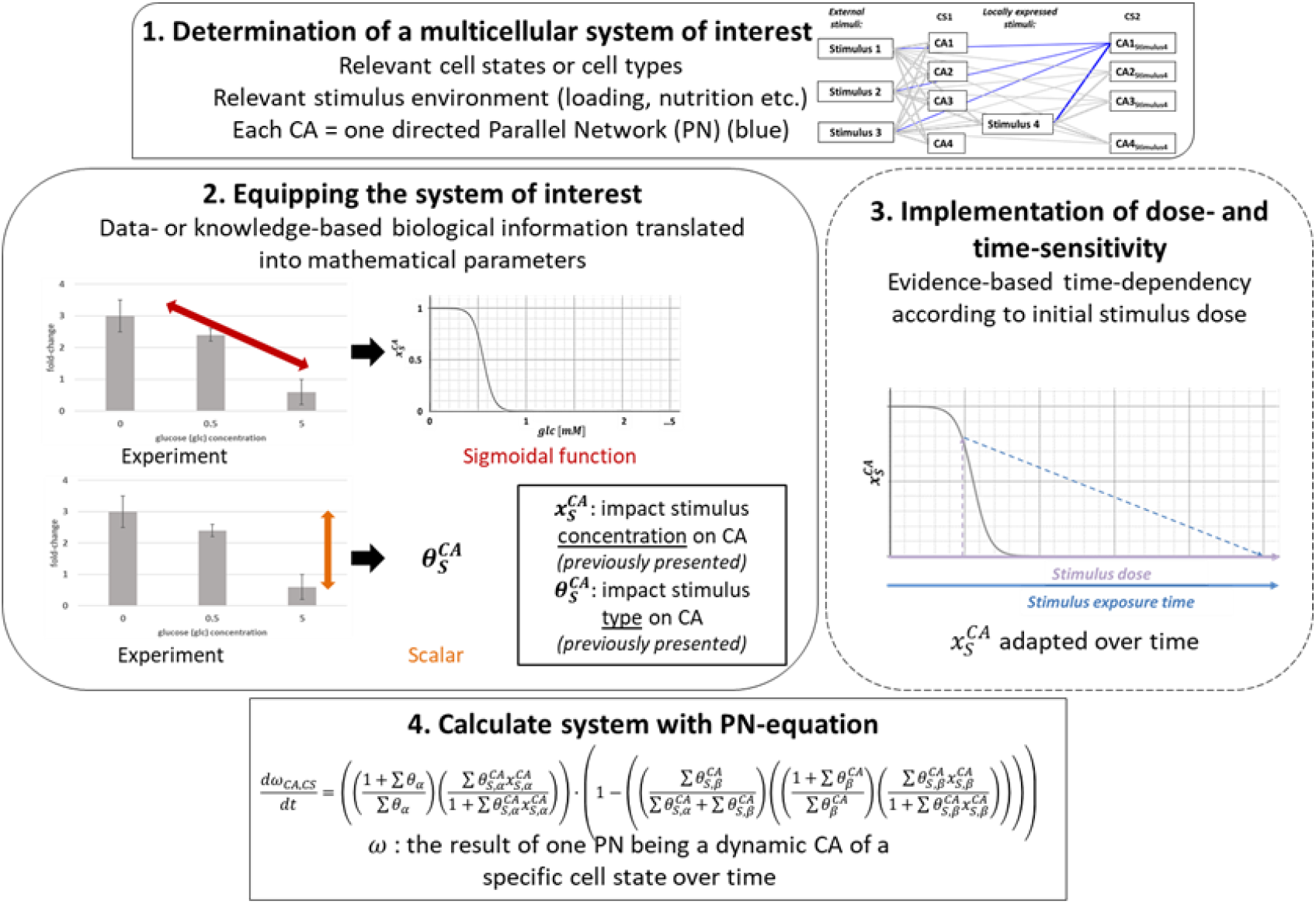
schematic workflow of the PN-Methodology, roughly divided into four principal steps (1. - 4.). 1. Conceptual, planning. (Exemplary network of 4 stimuli, 4 CA and 2 CS and one colored PN); 2./3. Mathematical methods to convert experimental information into mathematical parameters, whereby step 3 is not strictly necessary to apply the PN-Methodology; 4. First order ODE to calculate CA. CA: cell activity, CS: cell state.

In a first step (Step 1), a multicellular system of interest is determined. The system of interest identifies the targeted tissue (cell types and cell states), crucial (external and local) stimuli and CA (often mRNA expressions or protein synthesis). Step 1 is a conceptual, knowledge-based procedure, which results in a directed network, where stimuli are linked with the CA. Subsequently, each interaction within the directed network is equipped with mathematical parameters derived from experimental studies (17) (Step 2), and, if required, modulated to respond to time effects (Step 3). Eventually, the network is calculated with a specifically developed first order ODE, the PN-Equation (Step 4). In the next Sections, each Step (1-4) is explained in detail, and application is shown through the IVD case study. The basic terminology of the PN-Methodology is summarized in Table 1.

**Table 1:** overview over the basic terminology of the PN-Methodology.

| Terminology | Explanation |
| --- | --- |
| Cell activity (CA) | Cell responses (according to research interest / research question), e.g. in terms of mRNA expression or protein synthesis. |
| External stimuli | Stimuli (S) coming from an external source that globally regulate targeted CA. Such stimuli might be mechanical loads, nutritional factors, radiation etc. |
| Locally expressed stimuli | Stimuli that are locally expressed by cells within the simulated multicellular system. The chosen locally expressed stimuli are typically known for their ability to alter a targeted CA. |
| Cell state (CS) | Refers to a subgroup of cells (within a population of cells of the same type) that are exposed to an altered local stimulus environment due to locally expressed stimuli. |
| System of interest | Complete network that a user would like to investigate, consisting of all the CA, all the different CS and all the external and locally expressed stimuli. |
| Parallel network (PN) | Directed network that describes one CA, i.e. a specific mRNA expression or protein synthesis. Thus, one PN includes the tackled CA and all the stimuli that influence that CA. |
| PN-Equation | Specifically designed ODE that solves a PN. |
| PN-System | All the PN (CA) within the system of interest, where the user wants to obtain comparable results (profiles of different CA) |
| $x_S^{CA}$ | Sensitivity of a CA to a stimulus (S) concentration (17,18) |
| $\theta_S^{CA}$ | Sensitivity of a CA to a stimulus type (17) |
| $\omega_{CA,CS}$ | Result of the PN-Equation, i.e. a final CA of a given CS within the system of interest. |

### 2.2 System of interest and PN-Systems (Step 1, Figure 2)

#### 2.2.1 Determination of the system of interest

A system of interest consists of cell responses in terms of specific CA and of the stimuli that regulate the targeted CA. Hence, a cell response can be reflected as a profile of *n* targeted CA, regulated by a multifactorial environment of *m* stimuli. Such stimuli can either come from another source (external stimuli) or they can be locally expressed by the cells. Locally expressed stimuli cause *p* different cell states (CS), i.e., CS*p, p=1,II,III, …, p*, that co-exist within a multicellular environment. Stimuli can have either an activating, or an inhibiting effect on a CA. This activating or inhibitory nature can be predefined and fixed, or it can depend on the dose and the duration of a stimulus, i.e. overcoming simple on-off effects (Figure 3, top).

**Figure 3:**
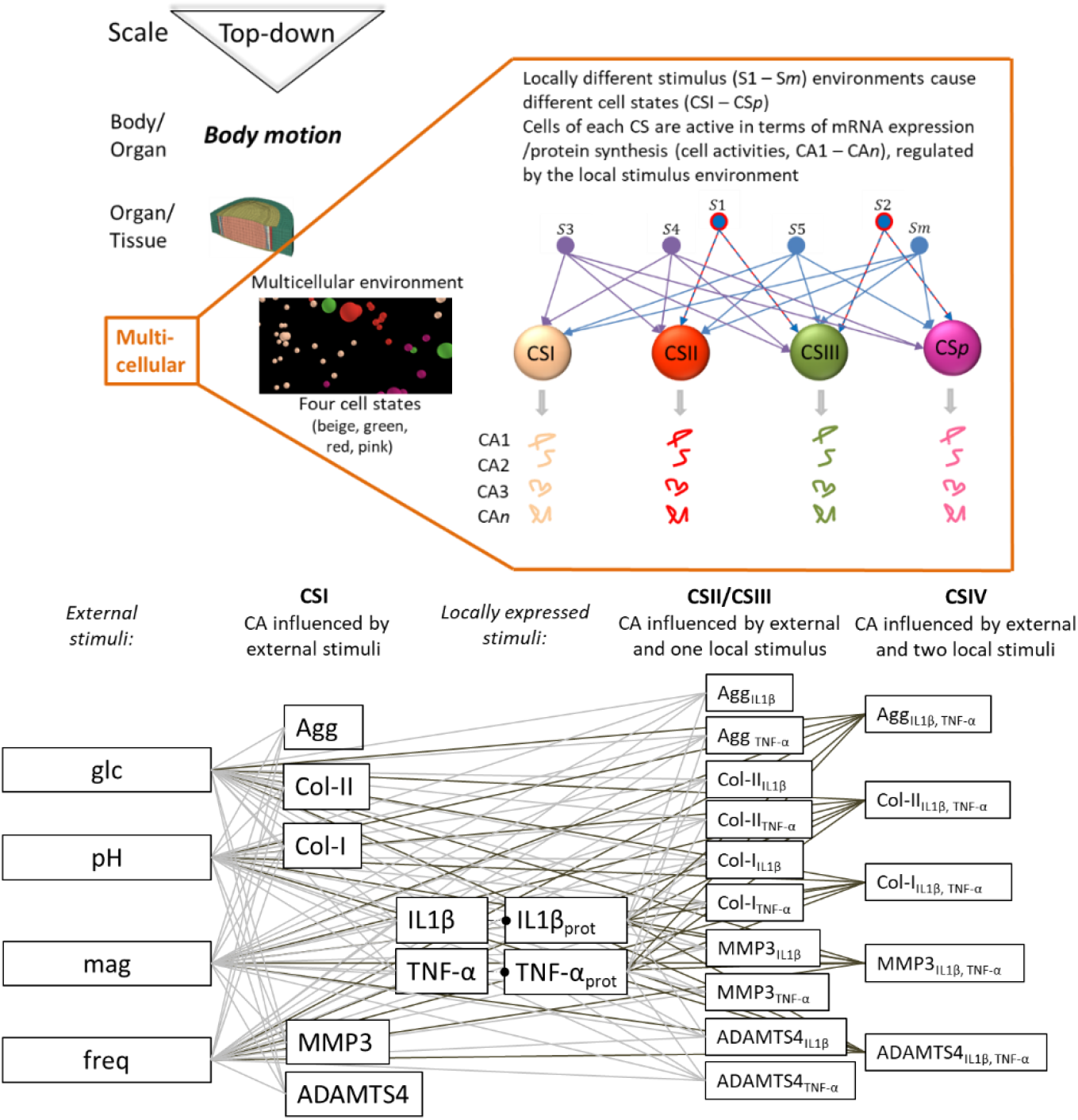
Top: the PN-Methodology tackles the multicellular level. Schematic visualization of a complex, multifactorial, multicellular environment, consisting of *m* stimuli and *p* cell states (CS) due to different stimulus combinations. *n* cell activities (CA) are analyzed for each CS. Individual stimulus-CA interactions (arrows) might have an activating (blue; *S1, S2, S5, Sm, Sn*), inhibiting (red; *S1, S2*) or a dose and time-dependent (purple; *S3, S4*) effect on a CA. Below: System of interest to approximate NP cell behavior, consisting of four external key stimuli (glc, pH, magnitude (mag), frequency (freq)). CA enclose the mRNA expression genes coding for tissue proteins (Agg, Col-I, Col-II) and proteases (MMP3, ADAMTS4). Different cell states (CSI-IV) were induced by altered stimulus environments due to the presence of the locally expressed proinflammatory cytokines TNF-α and IL1β. Prot: protein

In the case of the IVD NP, the CA of interest were the mRNA expressions of the main NP tissue components, Aggrecan (Agg), and Collagen Type II (4) (Col-II). Additionally, Collagen Type I (Col-I) was considered, as it becomes important in degenerative changes of the IVD (4). As per the protease-related CA, MMP3 and ADAMTS4, respectively from the matrix metalloprotease (MMP) and from the “a disintegrin and metalloproteinase with thrombospondin motifs” (ADAMTS) families, were considered, since these proteases are important in IVD degeneration (4,19,20).

Key relevant stimuli for the targeted CA were identified to be the nutrient-related factors, glc and pH, and the mechanical load parameters, magnitude (mag) and frequency (freq) (8,10–15,19,21). The relevant locally expressed stimuli were the proinflammatory cytokines interleukin 1β (IL1β) and the tumor necrosis factor α (TNF-α). IL1β and TNF-α were classically related to initiations of IVD degeneration (22), as they importantly alter the targeted CA of NP cells (23,24). In the multicellular environment of the NP, local expressions of proinflammatory cytokines cause heterogeneous cell environments, as NP cells have different CS, according to immunopositivity. Thereby, we were interested in i) how each proinflammatory cytokine individually affects the CA and ii) how the presence of both proinflammatory cytokines might affect a CA, compared to cells that are not exposed to proinflammatory cytokines. Hence, the PN-Methodology was required to predict the CA of four different CS: CA of cells only exposed to external stimuli (CSI), cells additionally exposed to one type of proinflammatory cytokine, i.e. either IL1β (CSII) or TNF-α (CSIII), and cells exposed to both types of proinflammatory cytokines, i.e. IL1β and TNF-α (CSIV). This eventually led to a system of interest for the NP as shown in Figure 3. From a mathematical point of view, the determination of the CS preserves the biological meaning and helps to properly define the PN-Equations (see Step 4, “Calculating PN-Systems with the PN-Equation”).

#### 2.2.2 Structuring the system of interest with relevant PN-Systems

PN-Systems are subsystems of the system of interest and must be defined prior to calculations, and before feeding the system with mathematical parameters. A PN-System contains all the CA, among which a relative, interrelated response, is required. For the current case study, an interrelated mRNA expression of tissue proteins and proteases of each CS is desired. Hence, all the CA of each CS belong to the same PN-System (PN-System I, Figure 4, A). In contrast, the prediction of locally expressed stimuli (proinflammatory cytokines) based on external stimulus concentrations, determines CSII, III and IV (Figure 4, A) and reflects individual PN-Systems (PN-Systems II&III, Figure 4, A). In this case, a proinflammatory submodel (previously reported (17)) estimates semi-quantitatively the level of proinflammatory cytokines used as local stimuli, which requires an estimation of corresponding proinflammatory cytokine mRNA expressions independently of the targeted CA. Examples for the mathematical handling of PN-Systems is given in Supplementary material 1.

**Figure 4:**
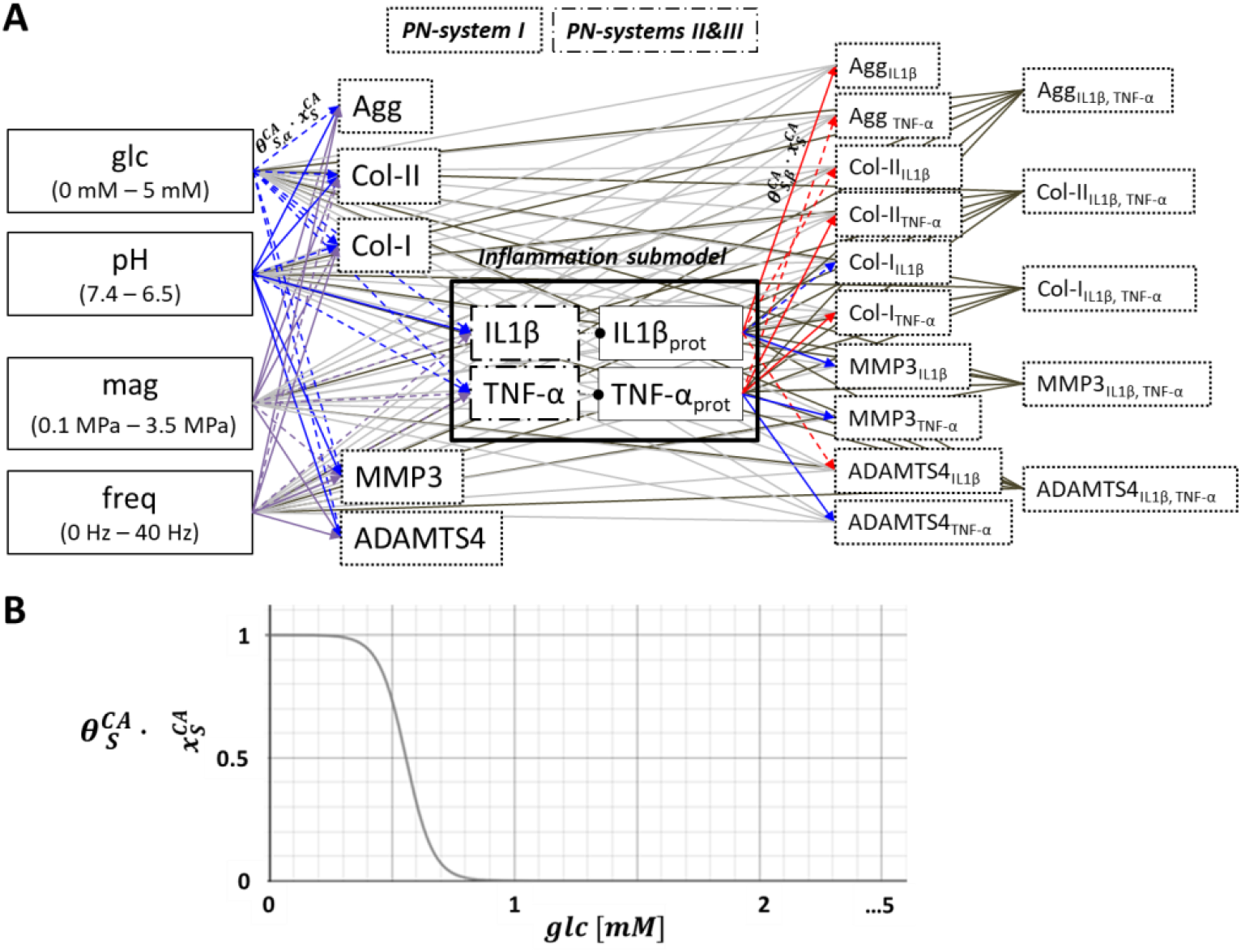
A: System of interest, subdivided into relevant PN-Systems. Arrows reflect the nature of individual stimulus-cell activity (S-CA) relationships, being either fixed, predefined as either activating (blue), inhibiting (red) or to be either activating or inhibiting depending on the dose of the dose of a stimulus (purple). Non-significant S-CA relationships are marked with dashed lines. The proinflammatory environment is estimated by an inflammation submodel (17). B: S-CA relationships are determined by a weighting factor 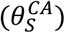 and a normalized sensitivity of a CA to a stimulus concentration 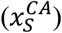. Example: glucose (glc)-MMP3 relationship.

### 2.3 Equipping the system of interest (Step 2, Figure 2)

The directed links within a system of interest that connect a stimulus to a CA are called S-CA relationships. S-CA relationships can be either generally activating, generally inhibiting or depending on the stimulus dose (Figure 4, A). For the current network, the activating or inhibiting effect of nutrients (glc, pH) and of proinflammatory cytokines were assumed to be predetermined and fixed (17): the generally present and less fluctuating levels of glc and pH imposed by the vasculature peripheral to the IVD along with indirect mechano-transduction phenomena (25) were used to obtain baseline levels of activation for each CA. Hence, nature of glc and pH stimuli was considered as always activating. In contrast, proinflammatory cytokines were integrated as either activating or inhibiting (Figure 4, A) based on experimental data from cell cultures (17). The impact of strongly fluctuating mechanical parameters on CA is assumed to be dose-and time dependent. While the handling of generally activating and inhibiting interactions was previously explained (17,18) and subsequently summarized, a method to approximate dose-dependency is introduced in section 2.3.1. S-CA relationships are determined by two variables, both directly derived from experimental research (Figure 2, Step 2; Figure 4, B). The sensitivity of a CA to a stimulus (subscript S) dose is reflected as 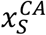 and is derived from the change of a CA due to the variation of a stimulus dose (18). The effect of the stimulus type was defined through a weighting factor 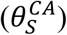 (17). 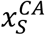 can obtain values between 0 (dose at which minimal CA was experimentally found) and 1 (dose at which maximal CA was experimentally found). The value of 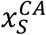 was approximated by one or various continuous sigmoidal functions, in order to relate each stimulus dose to a corresponding value for 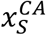(Figure 4, B). Biologically, the weighting factor, 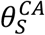, expresses the maximal perturbation of a CA provoked by a change of the stimulus concentration (ideally within a physiological range). Weighting factors are scalars that can take values of 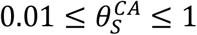, where the lowest weighting factor of 0.01 was assigned to non-significant changes of a CA due to varying stimulus concentrations (Figure 4, A, marked as dashed lines).

#### 2.3.1 Approximation of a dose-dependent activation or inhibition of a link

The effects of mag and freq on NP cells were simulated to be dose-dependent (Figure 4, A). First, relevant stimulus ranges were defined. For mag (here referring to intradiscal pressures) a relevant range from 0.1 MPa to 3.5 MPa was determined based on *in vivo* studies (26,27). The range for freq was set to 0 Hz – 40 Hz, aiming to allow predictions for the effect of whole-body exposures to high-freq in daily occasions.

Second, CA was estimated based on combined information from several experimental *in vivo* and *in vitro* studies (8,26–31), throughout the physiologically relevant ranges of mag and freq. As per mag, low and moderate pressures due to e.g. lying or walking were assumed to be highly anabolic. With rising intradiscal pressures, as obtained during jogging, anabolism lowers until the stimulus becomes catabolic. Additionally, we considered findings from Chan, Ferguson, and Gantenbein-Ritter (8) that suggest that mag around 0.2 MPa and 0.8 MPa reflect a potentially beneficial range of mag, whereas a (static) compression loading of >1 MPa induced degenerative changes. To estimate CA in function of freq, it was assumed that walking frequencies are overall beneficial. Walking freq might include freq of up to around 2.5 Hz as measured by Pachi et al. (29). In contrast, freq of 3-8 Hz, or over a threshold of 5 Hz, possibly disrupt proper cell function (31).

A generic behavior of anabolic and catabolic CA was initially assumed, and cell responses to mag and freq were represented through two generic (mirrored) functions for each stimulus, to reflect overall anabolic or catabolic behavior (**Figure 5**). The functions were fitted to an accuracy of four decimals, in agreement with the fitting of the continuous functions that describe the effect of nutrition-related stimuli (Figure 4, B, (17)). Generic functions evolving with rising stimulus intensities from activating to inhibiting were labelled with superscripts A;I 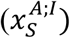 (**Figure 5**, top) and those evolving from inhibiting to activating were labelled with superscripts I;A 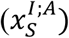 (**Figure 5**, bottom).

**Figure 5:**
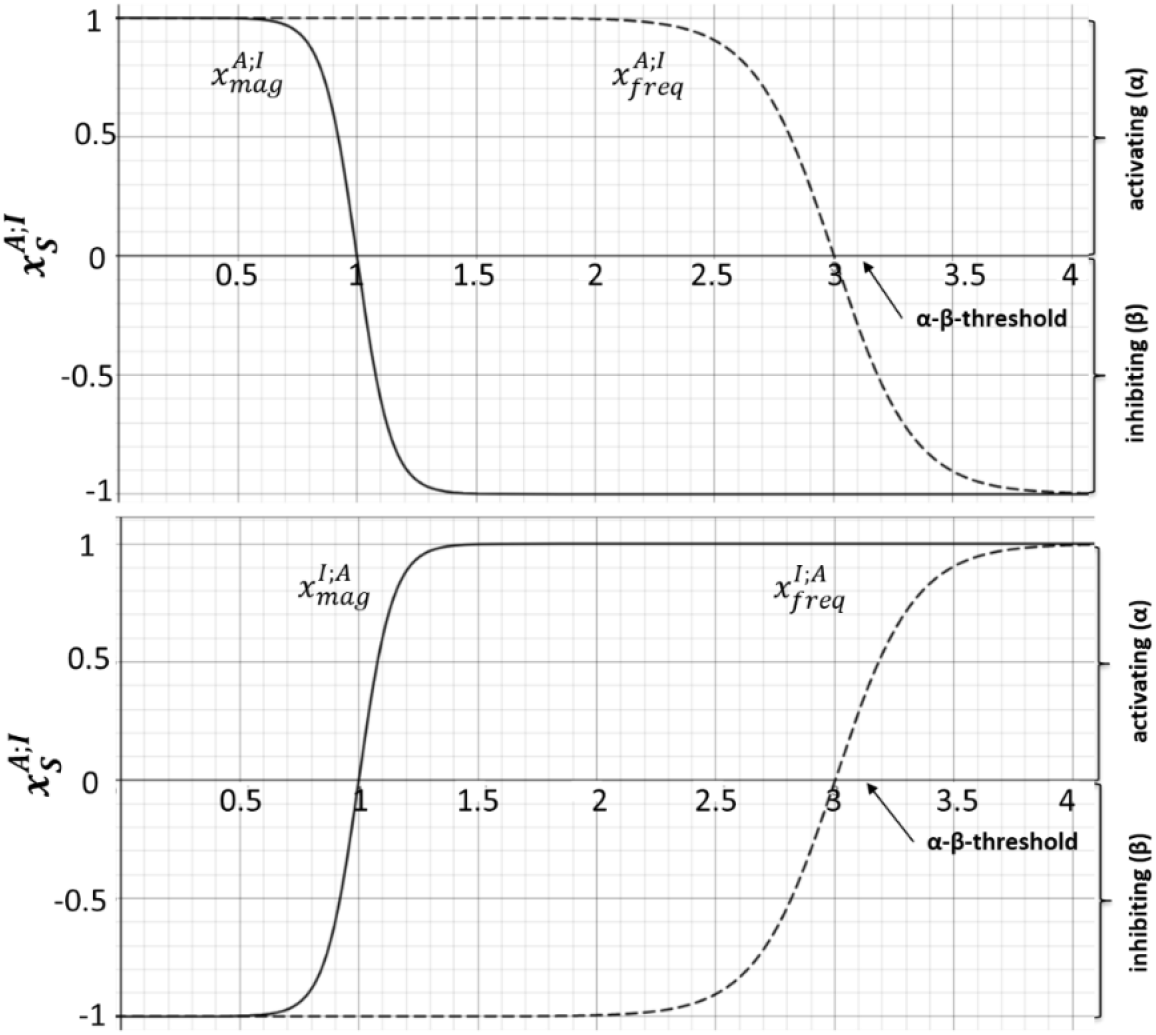
generic functions to approximate the effect of magnitude (mag,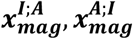) and frequency (freq, 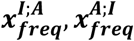)), respectively, on CA. Top: generic functions for CA that are assumed to be strongly activated under low mag turning to be strongly inhibiting under high mag (from activating to inhibiting A;I). Bottom: generic functions for CA that are assumed to be strongly inhibited under low mag turning to be strongly activated under high mag (from inhibiting to activating I;A)

The so-called α-β-threshold reflects the switch of a stimulus dose from activating (α) to inhibiting (β), or vice-versa. The range of *x*_*S*_ lies within [-1;1], i.e., −1 ≤ *x*_*S*_ ≤ 1. Thereby, 0 < *x*_*S*_ ≤ 1 reflects an activating effect of the stimulus on a CA, and 0>*x*_*S*_ ≥ −1 reflects an (actively) inhibiting effect on an CA.

The mathematical formulations of the four generic functions are given in Eqs. 1-4.

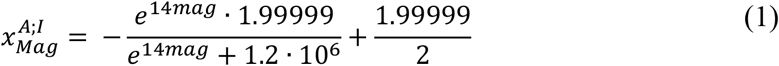

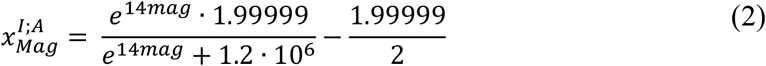

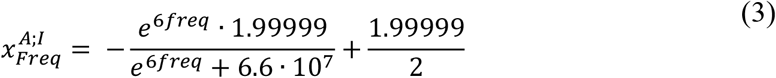

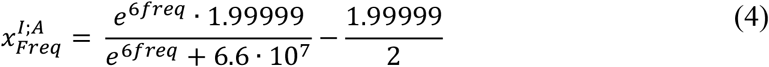

Agg and Col-II mRNA expressions were associated to the functions 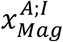 and 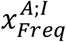, respectively, whilst any other mRNA expression was associated to 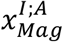 and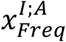.

Weighting factors 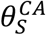 for mag and freq were obtained from an experimental study (15), where rat NP cells were exposed to a mag of 1 MPa and three different frequencies of 0.01 Hz, 0.2 Hz and 1 Hz. Weighting factors of mag were estimated using data of quasi-static conditions (i.e. 0.01 Hz). Weighting factors of freq were obtained based on the maximal x-fold change in mRNA expression caused by the three aforementioned frequency values (Table 2).

**Table 2:** individual weighting factors for loading parameters.

| Stimulus | mRNA | Weighting factor<br>($\theta_s^{CA}$) | Experimental study |
| --- | --- | --- | --- |
| Mag | Agg | 0.3484 | (15) |
|  | Col-I | 0.8711 | (15) |
|  | Col-II | 0.1394 | (15) |
|  | MMP3 | 0.0100 | (15) |
|  | ADAMTS4 | 0.1394 | (15) |
| | TNF-α | 0.0100 | Indications (32,33) |
| | IL1β | 0.0100 | Indications (34) |
| Freq | Agg | 0.3318 | (15) |
|  | Col-I | 0.4355 | (15) |
|  | Col-II | 0.0100 | (15) |
|  | MMP3 | 0.5124 | (15) |
|  | ADAMTS4 | 0.1048 | (15) |
| | TNF-α | 0.0100 | No information found |
| | IL1β | 0.0100 | No information found |

According to experimental measurements (32–34), the effect of mag on proinflammatory cytokine expression seems to be limited and information about the respective effect of freq was not found. Therefore, the weighting factors of TNF-α and IL1β was initally set to 0.01.

To obtain numerical values for weighting factors, the experimentally found maximal x-fold change in mRNA expression, was converted into a “cellular effort” as previously explained (17). The “cellular effort” allows to compare increases (1 < *ϵ* < ∞) and decreases (0 < *ϵ* < 1) of mRNA expressions with respect to a control level (1). A complete list of all weighting factors, together with x-fold changes and corresponding “cellular efforts” used in this study is provided in the Supplementary material 2.

### 2.4. Calculating PN-Systems with the PN-Equation (Step 4, Figure 2)

The PN-Equation is an ODE-based approach to resolve PN-Systems (Eq. 5). It was developed based on a well-established, graph-based analytical methodology to approximate single network dynamics (35) (Supplementary material 3). It determines a time-dependent overall activation of a CA of each CS 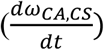 and is made of an activating and an inhibiting term (subscripts α and β, respectively):

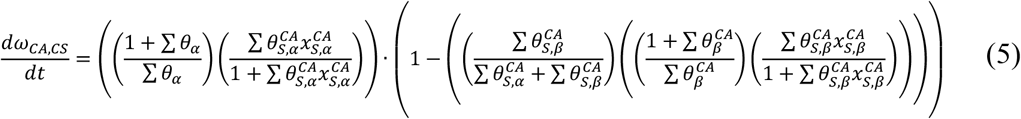

- ***Activation:*** The activating part of the PN-Equation (Eq. 5) determines an overall activation of a CA relative to any other CA within the same PN-System. Within the first term 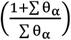, all activating weighting factors of the tackled PN-System were considered, which determines the size of the PN-System^1^. Hence, the maximal activation of one PN reflects a fraction of the overall size of the PN-System. The second term 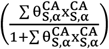 then defines the activation of the tackled CA, i.e. the activating component of a PN.
- ***Inhibition:*** The inhibiting part of the PN-Equation (Eq. 5) lowers the activation of a CA determined by the activating portion of the PN-Equation. It is defined relative to the activation of the CA. The first term 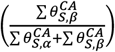 determines the relative weight of the inhibition on the tackled CA. Hence, if the activation of a CA was determined based on weak activators and powerful inhibitors, the impact of inhibition was strong and vice-versa. Importantly, this term prevents weak inhibitors to completely erase overall cell activations. The second inhibiting term 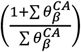 defines the maximal inhibition of a CA by considering all the different CS of a CA. This allows for coherent inhibiting impacts of different CS: its function is equivalent to the first activating term of the PN-Equation, but instead of considering the whole PN-System, only different CS of the same CA were taken into account. The third term 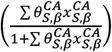 determines the actual weight of the inhibition of the tackled CA, analogue to the second term of the activating portion of the PN-Equation.

Figure **6** illustrates the determination of the inhibiting part of the PN-Equation. Further examples for PN-Equations for selected CA are provided in Supplementary Material 1.

**Figure 6:**
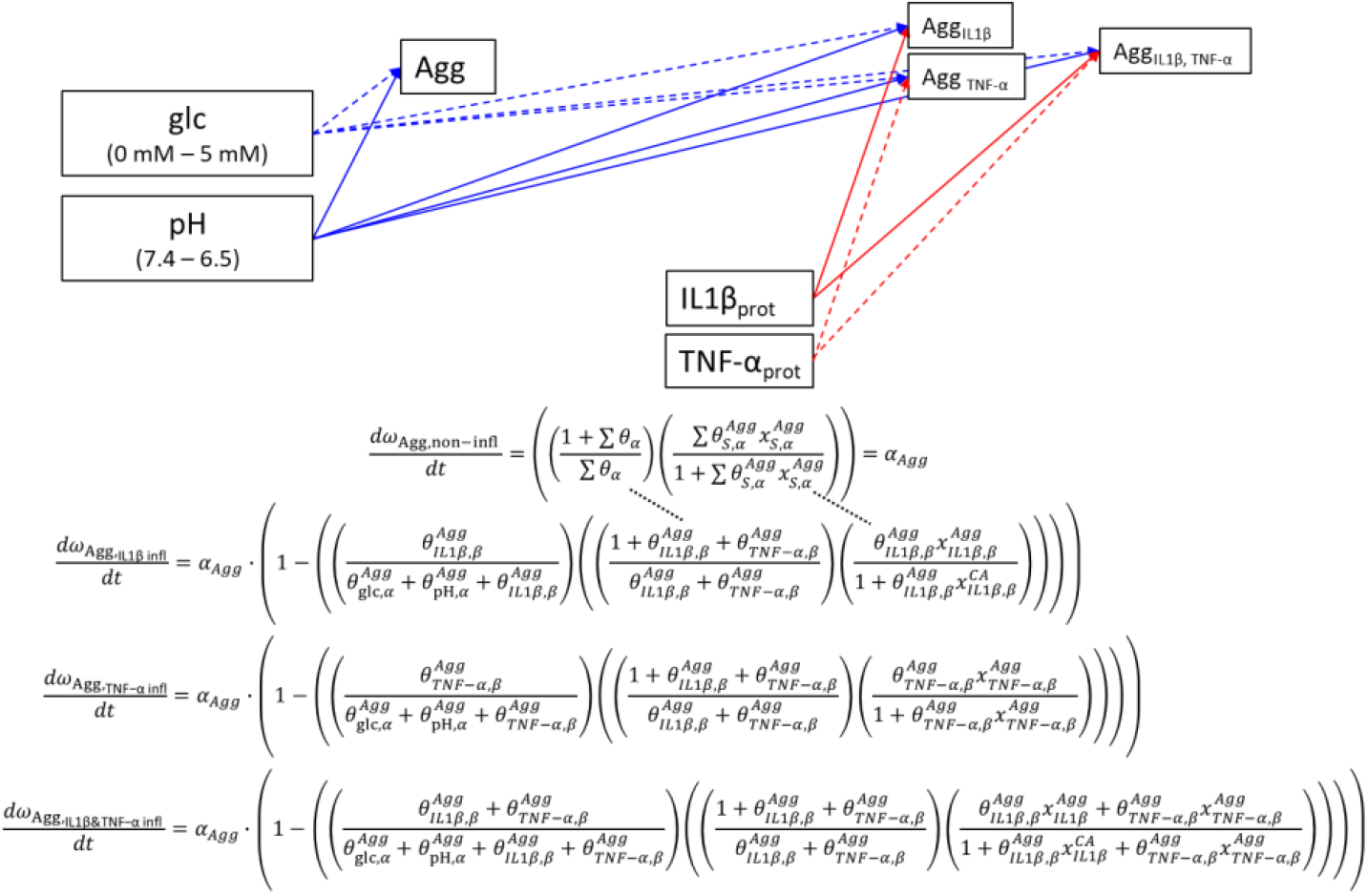
Illustration of the determination of the inhibiting part of the PN-Equation (here, the inhibition is caused by the impact of IL1β and TNF-α proteins (prot), respectively). Example for the different CS of Aggrecan (Agg) exposed under the influence of glc and pH. Analogue formulations between terms in the activating and inhibiting portions of the PN-Equation were indicated with dotted lines. Infl: inflamed

The result of the PN-Equation reflects the CA over time 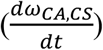 and is calculated as a dimensionless PN-activity. PN-activities semi-quantitatively reflect the relative activations of different CA within the same PN-System and actual, numerical values do not have a direct biological interpretation. The higher a PN-activity, the higher the tackled CA. PN-activities have a resolution of 4 decimals (17). The specific range over which an individual CA can vary is determined by the weighting factors of that CA. Hence, the more impact a CA has within the PN-System, the higher the weighting factors for this CA and the wider the range of values that this CA can obtain. If the user wants to know the maximal PN-activity for a CA, 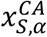 must be set to 1 and the inhibiting portion of the PN-Equation must be 0.

To compare the CA associated with different physical activities after a given time, the accumulation of PN-activities (acc. PN-activity) can be calculated, being the sum of a PN-activity over the given time.

### 2.5 Approximation of time-dependency (Step 3, Figure 2)

Time-dependency was integrated by introducing a time-sensitivity term (*τ*) in the 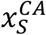 variable (Table 3). *τ* depends on three variables: the initial stimulus dose, a CA-specific time sensitivity 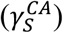 and the stimulus exposure time (t). For the IVD case study, mag and freq effects were time-sensitive (8,16,36), and the knowledge-based generic functions presented as Eq. 1-4 (section 2.3.1.) were coupled to *τ*. Therefore, Eq. 1-4 were re-introduced as *X*_*S*_ (Eq. 6-9), out of which *x*_*S*_ will eventually be calculated (Table 3):

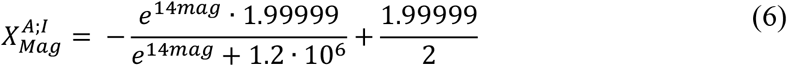

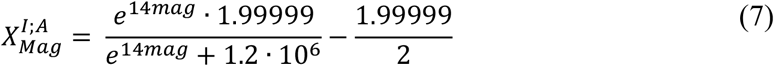

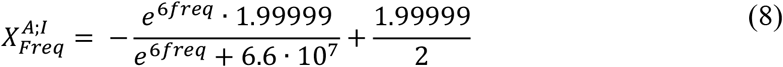

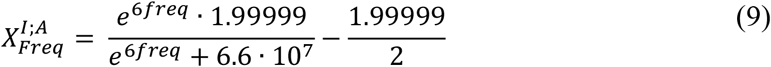

*X*_*S*_ defines the initial stimulus dose, based on which the effect of time was calculated. Hence, it determines the velocity at which the anabolic effect of a stimulus decreases, or at which the catabolic effect increases. It was either obtained through the absolute difference between 1 and 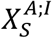 and between 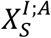 and -1, or through the (absolute) value of *X*_*S*_ (Figure 7, Table 3).

**Table 3:**
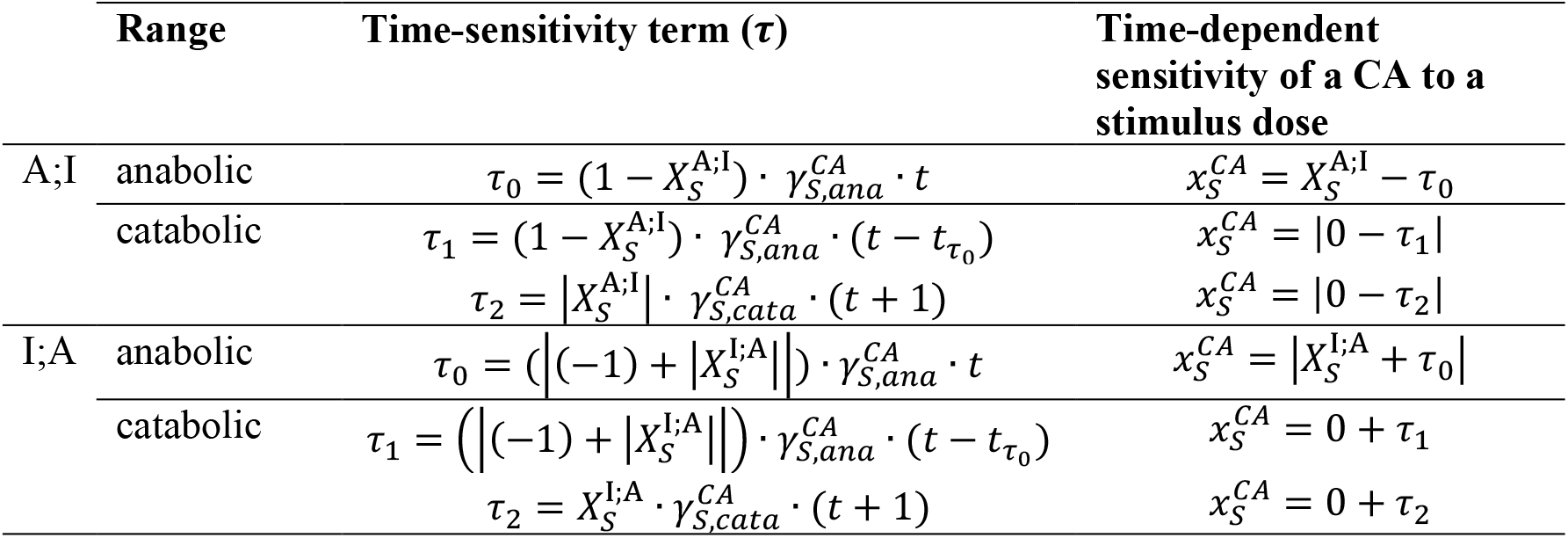
mathematical description of the time sensitivity term (***τ***) and the sensitivity of CA to a stimulus dose 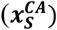. Note that integration into the PN-Equation requires positive values. Hence, absolute values were used for the terms known to be negative. A;I: generic functions that evolve from activating to inhibiting. I;A: generic functions that evolve from inhibiting to activating. ***τ*** are schematically plotted in Figure 8, a (A;I) and b (I;A), respectively.

**Figure 7:**
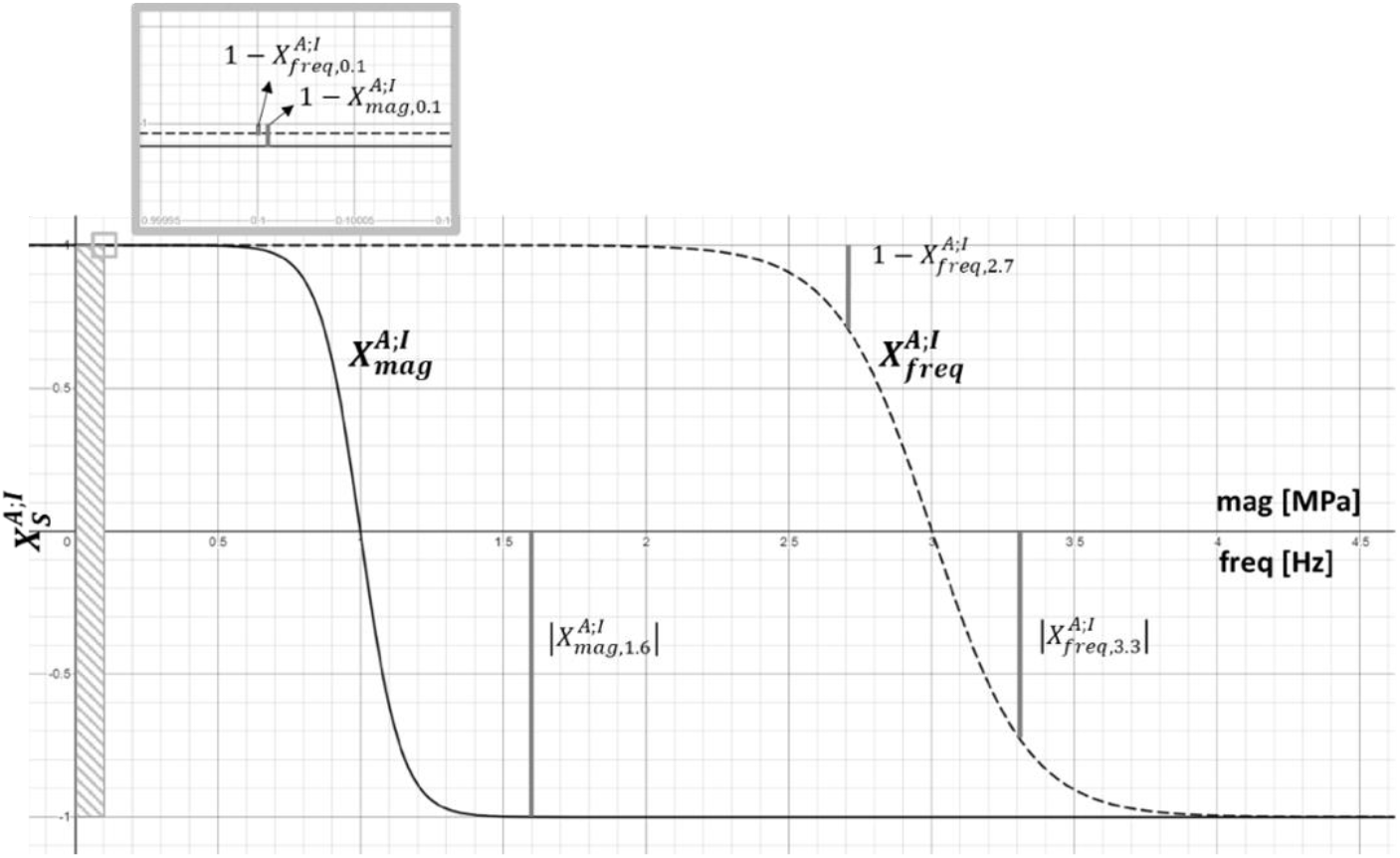
determination of the influence of a stimulus dose on the velocity at which the anabolic effect of the stimulus gets lost or at which the catabolic effect reaches its whole potential. Example for 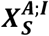 functions of mag and freq. Illustrations for the stimulus intensities at 0.1 MPa, 1.6 MPa and 0.1 Hz, 2.7 Hz and 3 Hz. Lowest values for mag reflect 0.1 MPa.

The overall CA-specific time sensitivity, 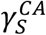, consists of the product of two parameters: an adjustment factor (*σ*), to translate numerically the range of values generated by the generic functions into a biologically relevant range and an individual (subscript i) time sensitivity of each CA derived from experimental data 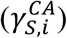 (Eq. 10).

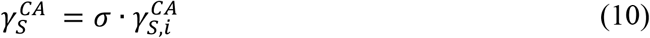

An approximation of *σ* and details to obtain CA-specific time sensitivity 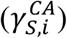 for the IVD case study are provided in Supplementary material 4 and 5. The mathematical formulation of *τ* varies depending on the initial stimulus dose (Table 3).

*τ*_0_ defines the loss of the effect of an initially anabolic stimulus over time. After passing the α-β-threshold (i.e. when an initially activating stimulus becomes inhibiting due to prolonged exposure time) *τ*_1_ is activated. *τ*_1_ has the same mathematical description as *τ*_0_, but considers as a time origin the individual time point 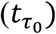, when the α-β-threshold was crossed. *τ*_2_ describes a time delay for a catabolic stimulus to reach its whole catabolic potential. The effect of time sensitivity is applied until a maximal activating 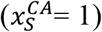 or inhibiting 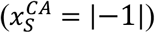 impact of a stimulus is reached (Figure 8).

**Figure 8:**
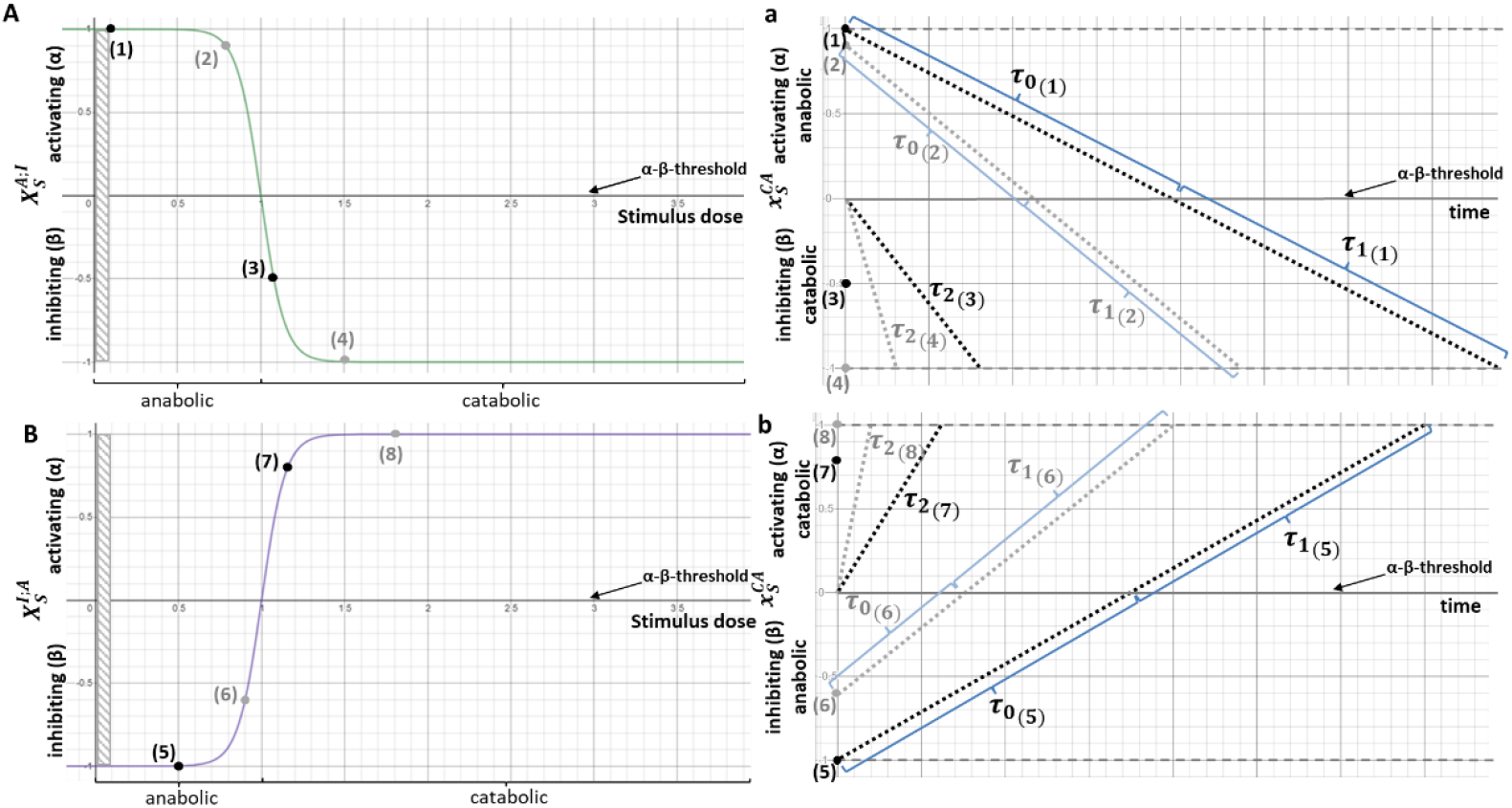
Illustration of mirrored generic functions 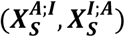, example of magnitude (see Figure 5) evolving from activating to inhibiting (A;I) (A) and from inhibiting to activating (I;A) (B) under rising stimulus intensities. a, b illustrate the time dependency (***τ***) that finally determines 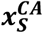. The influence of the stimulus dose on the time-sensitivity was schematically sketched for eight randomly chosen stimulus intensities ((1)-(8)); four on each mirrored generic function (A and B). Hence, the anabolic effect of the most anabolic stimulus dose ((1), A) remains over longer time periods ((1) a) than a less anabolic stimulus (e.g. (2) A, a). Time dependencies reflect a loss of activation due to a stimulus exposure over time (***τ***_**0**_), and a corresponding active inhibition after passing the α-β-threshold (***τ***_**1**_). For initially catabolic stimulus intensities, a latency time before developing a maximal negative impact on a CA was considered (***τ***_**2**_). Extremal 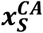 are 1 and -1, respectively.

#### 2.5.1 Approximation of static mechanical loading

As shown in Figure 7, current generic functions generally lead to a more anabolic response at lower doses compared to higher doses of mag and freq. Hence, a dose of mag = 0 MPa or freq = 0 Hz, would be treated more anabolic than any higher dose of load, as the value of the function is closer to 1 (or -1, respectively). While mag = 0 MPa is not considered for being out of the physiological range for the NP, a freq = 0 Hz represents static mechanical loads. Since static activities are generally less anabolic than moderately dynamic activities (8), the generic function was discontinuous towards freq = 0, i.e. between 0 < *freq* ≤ 0.1. Accordingly, freq = 0 Hz was described as a special case. To consider static loading, the impact of “no-frequency” on a CA must be estimated. Therefore, the information that degeneration might not be induced during 8 h dynamic axial loading with a load of 0.2 - 0.8 MPa and 0.1-1 Hz (8) was used. Hence, a dynamic daily human physical activity with loading parameters within the suggested range was compared to a generally anabolic, static, human behavior. As a dynamic human moving activity, comfortable walking was chosen. It was approximated by a mag of 0.60 MPa, reflecting a (rounded) average intradiscal pressure during walking (26), and a freq of 1.00 Hz. The static control was chosen to be “sleeping”, reflected by an intradiscal pressure of 0.15 MPa, being the (rounded) average intradiscal pressure during night, over a period of 7 h (26). Agg was chosen as the reference protein for the fitting (Figure 9).

**Figure 9:**
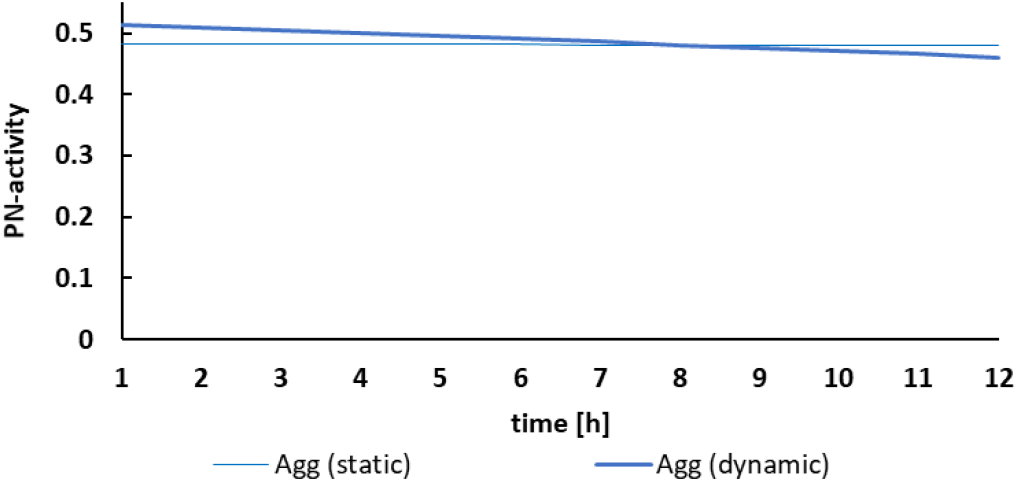
Illustration of the model prediction with ***X***_***freq***,**0**_ = **0. 75** for Agg (ref).

Results showed that a loss of anabolism due to dynamic loading compared to a static control after 8h was approximated with *X*_*freq*,0_ = 0.75. This means that 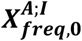 has a value of 0.75 instead of ≈1, whilst the coupled time-sensitivity (*τ*) for static loading was defined using the generic function of freq at 0 Hz (i.e. 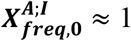 in Figure 7).

#### 2.5.2. Integration of time-dependent effects into the PN-Equation

As shown in section 2.3.1., the range of 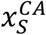 of mag and freq is bound between [-1;1]. Thereby, 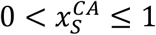 is integrated within the active portion 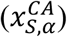 and 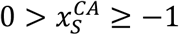 is integrated within the inhibiting portion 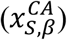 of the PN-Equation. Note that only the absolute values of 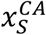 are then fed into the PN-Equation, as the latter only accepts positive values for 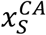 bound between 0 and 1 (see Supplementary Material 3). Hence, the sign provided by the logistic function indicates the part of the PN-Equation in which the value of 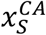 should take a role.

If a stimulus contribution changes from anabolic to catabolic due to time-dependency, the corresponding weighting factor, *θ*, must be listed in the two first terms of the inhibiting part of the PN-Equation (Eq. 5). Weighting factors within the term, where the stimulus is initially not active (including second term activating portion, third term inhibiting portion, respectively), is defined as “completive weighting factor” (indexed with *δ*) and can be understood as placeholder within the PN-Equation (see also Supplementary material 1). Completive weighting factors have the same value as their counterparts. Hence, if the weighting factor that describes the impact of mag on Agg equals 0.3484 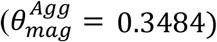 the corresponding, completive weighting factor, i.e. 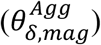 has the same value.

### 2.6 Evaluating the performance of the PN-Methodology with time-sensitive PN by calculating cell responses to microgravity and to various daily human activities

CA of each CS, i.e. immunonegative cells, cells immunopositive for either IL1β or TNF-α and cells immunopositive for both; IL1β and TNF-α, were calculated for optimal nutritional conditions (pH 7.1, 5 mM glc) (11,37). Two sets of simulations are provided; a first set of results encloses predictions for exposure to microgravity during six months (180 days) of space flight compared to a corresponding time spent on Earth. A second set of results was obtained by simulating cell responses of non-inflamed cells for eight human physical activities: sleeping, sitting with an active or a round back, hiking with or without extra weight of 20 kg, walking, jogging and sitting in/on a motor vehicle (Table 4).

**Table 4:** overview of the simulated human activities, corresponding loading parameters magnitude (mag) and frequency (freq), and duration (h).

| Human activity |  | mag<br>[MPa] | freq<br>[Hz] | time<br>[h] |
| --- | --- | --- | --- | --- |
| Microgravity |  | 0.25 | - | 4320 |
| Average day on Earth | Sleeping / laying down | 0.15 | - | 10 |
|  | Sitting with an active back | 0.55 | - | 2x4 |
|  | Walking | 0.60 | 1.80 | 3x2 |
| Daily physical activities |  |  |  |  |
|  | 1 Sleeping | 0.15 | - | 2;8 |
|  | 2 Sitting, active back | 0.55 | - | 2;8 |
|  | 3 Sitting, round back | 0.85 | - | 2;8 |
|  | 4 Hiking without extra weight | 0.60 | 1.00 | 2;8 |
|  | 5 Hiking, 20kg extra weight | 1.10 | 1.00 | 2;8 |
|  | 6 Walking | 0.60 | 1.80 | 2;8 |
|  | 7 Jogging | 0.65 | 2.75 | 2;8 |
|  | 8 Sitting in/on a motor vehicle | 0.55 | 15.00 | 2;8 |

Details regarding the estimation of intradiscal pressures and freq are explained in Supplementary material 6.

The current results were calculated with the Agent-based modeling software NetLogo (38), v. 6.0.2 (3D), as the calculated CA was coupled to a 3D NP cell environment (17). Based on the user-defined input data (nutrition concentrations, loading conditions and the exposure time (in hours, Table 4)), the CA of the multicellular environment is calculated and the current mRNA profiles for each CS is provided. At hour 0, values for mRNA expression are 0, establishing a value after 1h user-defined conditions. Results for the physical activities were provided both as a continuous evolution over 8 h and as an accumulation over 2 h.

## 3. Results

An approximated daily life on Earth led to high CA in terms of mRNA expression of the functional structural proteins of the tissue (Agg, Col-II) and to (close to) minimal CA in terms of protease mRNA expression in non-inflamed cells. Inflammation caused an overall yet modest upregulation of proteases and a slight downregulation of tissue proteins (Figure 10, A2).

**Figure 10:**
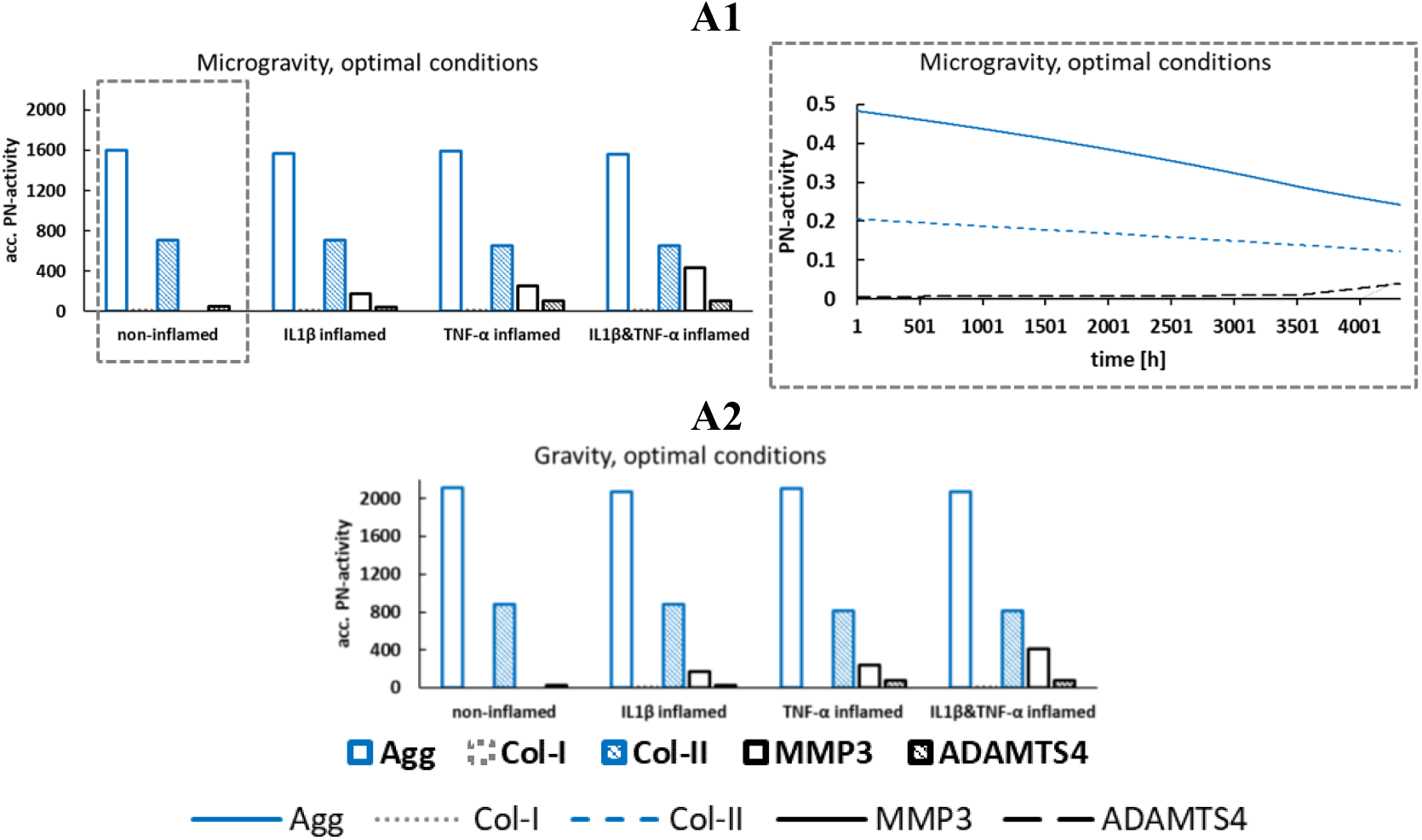
CA-profiles under microgravity: accumulated (acc.) PN-activity (A1, left) for four different proinflammatory cell states and the (continuous) PN-activity of non-inflamed cells (A1, right). CA-profiles of four different proinflammatory cell states (A2) for an approximated daily life on Earth. Time period: six months (180 days, 4320 h).

In contrast, exposure to microgravity (Figure 10, A1, left) led to an overall reduced PN-activity of the main tissue proteins to roughly 75% and 80% for Agg and Col-II mRNA expression, respectively. MMP3 mRNA expression was overall similar and ADAMTS4 was enhanced, though it remained low. The rise in ADAMTS4 protease expression could be especially attributed to the end of the stay in space (Figure 10, A1, right).

Regarding specific physical activities (Figure 11), high anabolism was found for sleeping (Figure 11, B1, B2), sitting with an active back (Figure 11, B1, B3), walking (Figure 11, B1, B4) and walking without extra weight (Figure 11, B1, B5). Catabolic shifts were predicted for any other condition after either 1h or after prolonged exposure. Especially hiking with heavy weights (Figure 11, B1, B6) and exposure to vibration (Figure 11, B1, B9) appears to be critical, given that a catabolic shift was already expected within the first hour of stimulus exposure.

**Figure 11:**
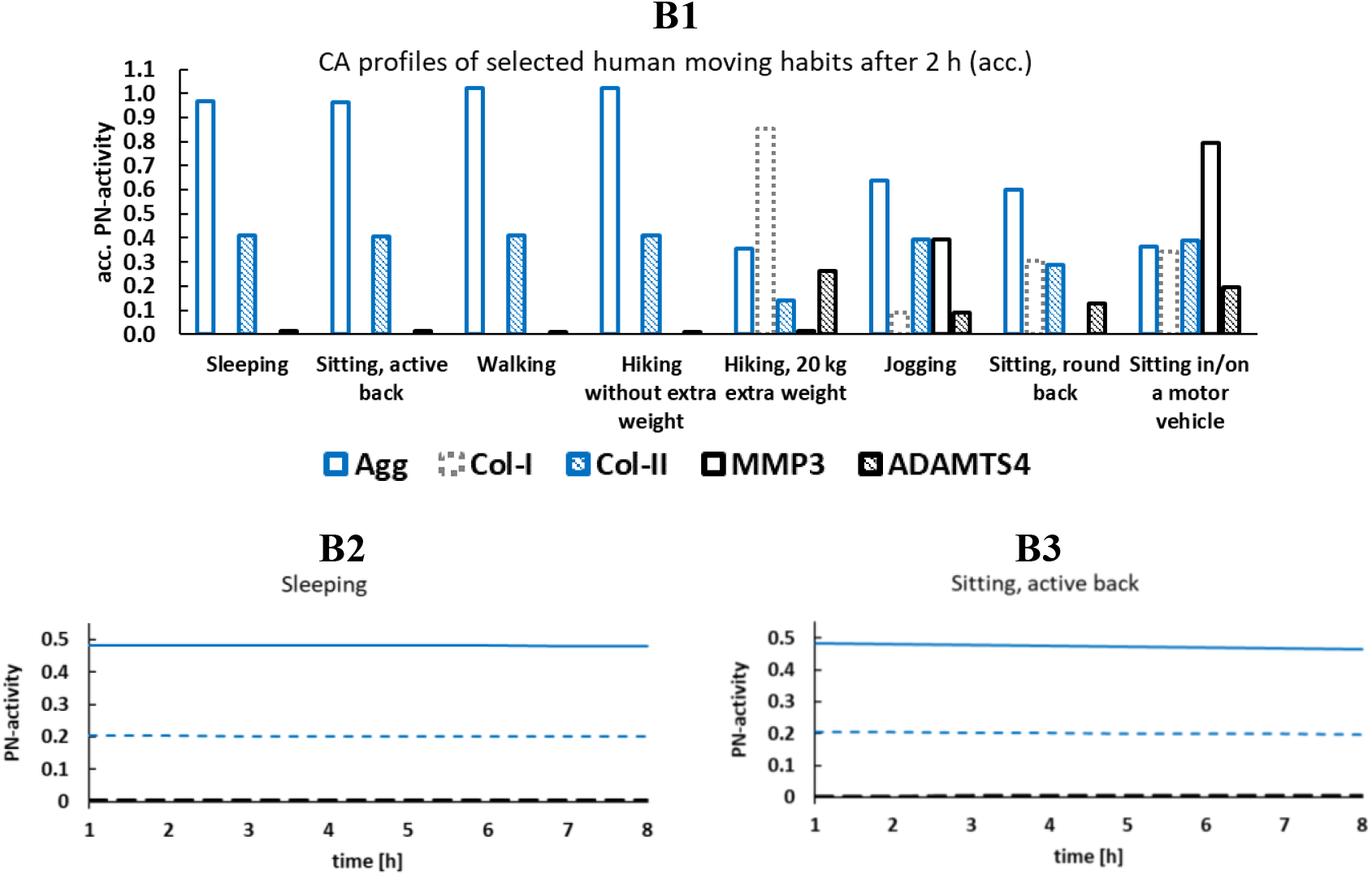
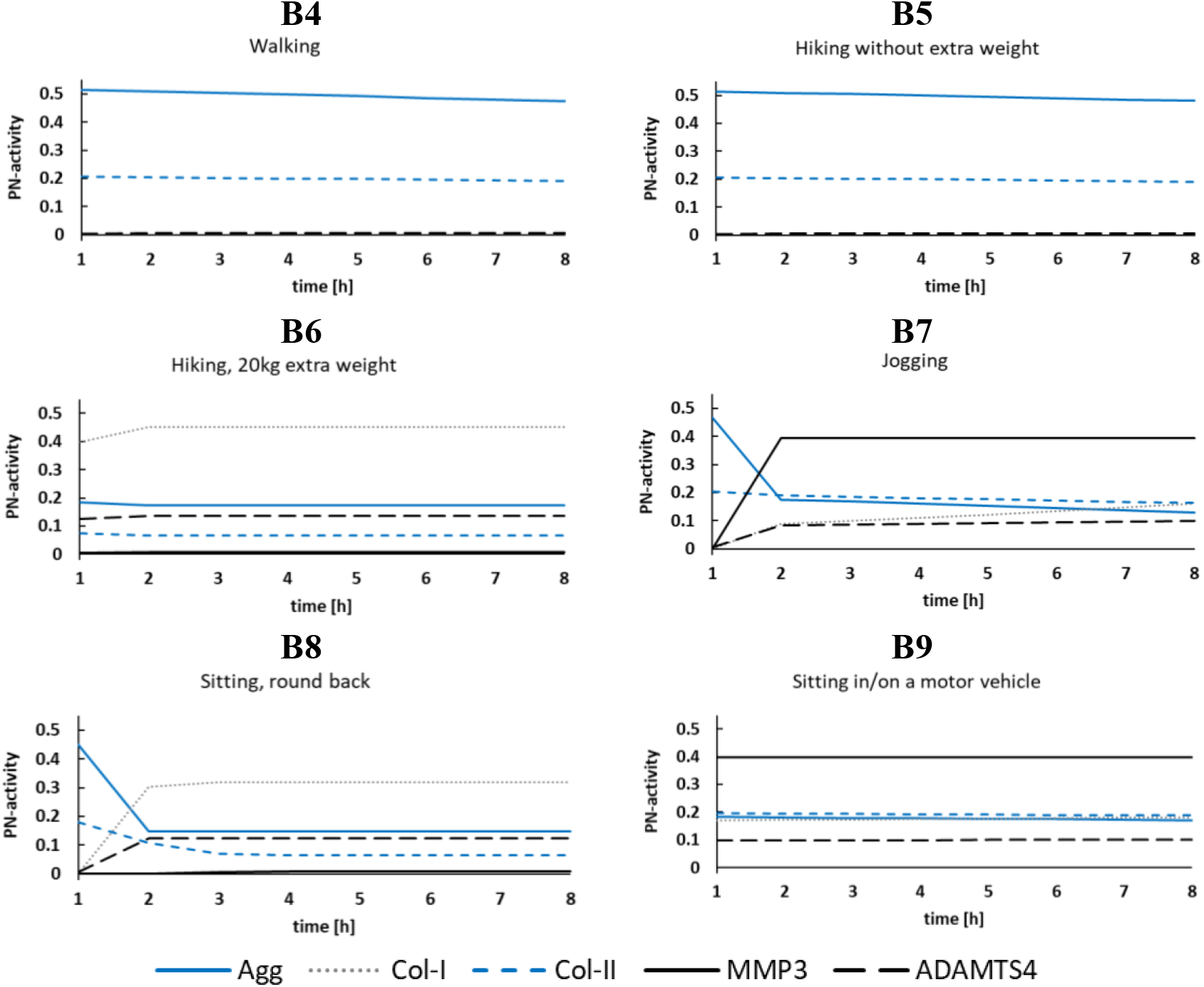
Predictions for eight selected physical activities: sleeping, walking, hiking with/without extra weight, jogging, sitting with an active/round back, exposure to vibration (as probably experienced by a motor vehicle). Accumulated (acc.) PN-activities after 2h (B1), and continuous PN-activities (B2-B9) over 8h of physical activity.

## 4. Discussion

The PN-Methodology is a novel deterministic network modeling approach that interprets dynamics in multicellular systems as interrelated parallel ongoing actions. It represents a feedforward high-level methodology to combine existing experimental knowledge to virtually estimate the responses of cells within multicellular systems of complex, multifactorial environments. The high-level component was introduced by linking directly critical external and local stimuli to relevant CA, therefore omitting the modeling of underlying intracellular regulatory networks. As a result, interrelated CA profiles for various CS were obtained for user-defined stimulus environments. Additionally, a method was developed to achieve time-sensitive cell responses, leading to a dynamic network modeling approach.

### 4.1 The PN-Methodology as a novel mathematical technique to calculate CA within heterogeneous, multicellular environments (Step 1, 2, 4, Figure 2)

The concept of parallel networks is, to our knowledge, a novel concept in network modeling of biological systems, complementing current approaches of mechanistic modeling (1,2). A major difference to knowledge-based graph representation is that this latter sets up topologies based on a priori connections that need to be verified and/or inversely calibrated (reduced or enriched) a posteriori. The PN-Methodology uses readily driven forward topologies defined by the experimental measurements that implicitly incorporate all hidden biological interactions. Hence, these interactions are incorporated in the measured S-CA relationships. Eventually, due to the high-level concept, the required experimental data to semi-quantitatively describe a biological system is reduced compared to bottom-up approaches aiming to simulate a system of similar complexity. Importantly, this modeling approach does not require links of a fixed (activating or inhibiting) stimulus nature. The model framework allows to change the stimulus nature, depending on the stimulus dose. This is not common for graph-based (logical-) modeling approaches (1), yet required, at least for (multi-) cellular systems, as it enables to represent reported biological sensitivities to varying stimulus doses (e.g., for mechanical loading, reviewed in (8)). A further advantage of this methodology is the concept of interrelated equation parameters through 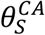 to obtain decoupled systems of ODE with exclusively analytically solvable ODE. Hence, the PN-Methodology can be calculated with any mathematical software able to resolve basic mathematical operations, such as the implementation of sigmoidal functions or the solving of ODEs. The methodology was specifically designed to work at a low computational cost, hence, on an “ordinary” personal computer, the presented results, including six months of microgravity exposure, were calculated within minutes.

The provided framework reflects a generic and scalable design, adaptable to user-defined systems of interest. This is a requirement for both, an extended investigation of the currently presented network, and the application of the PN-Methodology to multicellular systems of other tissues. Whilst the presented case study focussed on the CA of the same cell type, i.e., the NP cells, for different CS, the PN-Methodology would technically allow to model interrelated CA of more than one cell type and their corresponding CS, given that it does not rely on cell type-specific intracellular pathways. An integration of information is additionally facilitated, since the methodology allows the introduction of both data- (previously presented (17,18)) and knowledge-based information.

The PN-Methodology provides semi-quantitative results. Thereby, the range within which a CA can vary depends on the strength of the corresponding PN, which is defined through experimental information, i.e. through the weighting factor *theta*. Hence, ranges are highly CA-specific, rather than bound within a normalized range (i.e., between 0 and 1), as provided by logic-based (Boolean or Fuzzy Logic) network models (2).

The objective of the Methodology to virtually combine experimental data makes the system highly sensitive to the experimental input. Hence, a sensitivity analysis would confirm a high sensitivity to the input parameters defined by the experiments. Uncertainty is obviously directly derived from experimental data, i.e. affecting the *x* and *θ* parameters of the PN-Equations that describe the system. This is expressed through standard derivations of biological experiments, and by similar experimental setups, leading to differences in quantitative cell responses (e.g. (11,12)). In this article, however, focus was set on the introduction and evaluation of the novel mechanistic methodology, rather than on an investigation of the effect of uncertainty of experimental data on simulation results. Whilst strengths and limitations of *x* and *θ* parameters per-se were previously discussed (17,18), including limitations due to the use of animal experimental data or suboptimal cell culture conditions, the present discussion focusses on the PN-concept per-se.

Parallel networks are calculated with the PN-Equation, which was developed based on ODE designed to describe the interconnectivity of knowledge-based logical models (35). In this original approach and without integration of time sensitivity, ODE converge to a steady state, which is the output eventually analyzed.

The highly conclusive qualitative results obtained for the IVD case study were compared to numerous findings in the literature and holistically analyzed and discussed in our supplementary material 7 & 8. Moreover, a further study applying the PN-Methodology to another set of CA showed its adaptability to successfully predict CA profiles for other systems of interest (39).

Limitations of the PN-Methodology include that the PN-Equation currently only considers parallel effects of different stimuli, whilst possible cross-effects among stimuli were not taken into account. Yet, the equation allows to introduce correction factors for stimulus interactions, e.g. by formulating the weighting factors as variables, in function of the surrounding stimulus environment, if experimental evidence points towards significant impacts of cross-effects. Model dynamics by adding such an additional dimensionality remains to be investigated.

The PN-Equation favors increasing over decreasing PN-activities. Hence, tissue protein expressions decrease slowly, whilst the rise for proteases is more pronounced. Yet, good qualitative correspondence between independent literature information and current results (supplementary material 7 & 8) suggests that this tendency fits well the known biological dynamics in the NP of the IVD.

The simulation results of (multi-)cellular systems, as presented hereby, might be coupled to either lower scale models that represent intracellular regulatory networks able to further inform a CA, or to upper scale (FE) models where CA inform the biological regulation of homogenized model parameters (e.g., tissue composition parameters). Hence, the PN-Methodology has the potential to comprehensively bridge the gap between intracellular modelling and predictions at the tissue level.

### 4.2 Obtaining dynamic environments with time-sensitive PN (Step 3, Figure 2)

Exposure time is assumed to crucially influence cell responses within the IVD (8). An integration of time-sensitivity is therefore key to approximate dynamic responses of the multicellular system to chronic stimulus exposure over time. This considers a possible change of the nature of a stimulus (from activating to inhibiting or vice versa), due to both stimulus dose (see section 4.1), and exposure time. In contrast, state-of-the-art deterministic graph-based approaches (e.g. (35)) are usually not able to consider the effect of exposure time of individual links (1,35). Hence, integration of the time effect, combined with a mathematical solution to control the nature of the stimulus link, is a unique asset of the PN-Methodology to approximate semi-qualitatively the innate dynamics of biological systems. Arguably, numerical incongruencies close to the transition of a stimulus between activating and inhibiting, i.e. around the α-β-threshold (Figure 8), occur due to the use of continuous functions, which is extensively discussed in supplementary material 9. Yet, current simulations allow comprehensive predictions of the dynamic regulations of the system (Figure 10, A1, right, Figure 11), the biological context and relevance being also extensively discussed in supplementary material 7. Time-dependencies were programmed to be gradual and linear and became non-linear due to integration into the PN-Equation. Despite the overall good qualitative agreement of our results with physiological findings (supplementary materials 7 & 8), the effect of time might be non-linear. The model can be adapted accordingly, as soon as further insights into the time-responses of NP cells become available, by reconsidering the time variable as a function. Moreover, the straightforward design of the mathematical framework allows an easy replacement of any generic function if data-based information become available. Currently, time dependencies were integrated for mechanical stimuli because of their well-known dose- and time dependent effect on NP cells (8). However, technically, they can also be integrated for biochemical stimuli.

Thanks to the integration of time-dependent PN into the PN framework, the system continuously evolves over physiologically relevant periods of time. Hence, even though some simulations appear to reach a steady state at first sight, e.g. sleeping (B2) or sitting with a round back (B8) (Figure 11), PN-activities are constantly evolving over long periods of time, which can be visualised by prolonging the simulated time period: sleeping shows similar dynamics as exposure to microgravity (Figure 10, A1 left), due to the similar mechanical conditions (Table 4). Dynamics might be also observed in different CS. This is illustrated in Figure 12, where sitting with a round back is simulated for cells immunopositive for IL1β and TNF-α, over 8 h. The same conditions with non-inflamed cells lead to different and more beneficial regulation of CA (Figure 11, B8). This analysis of the effect of CS shall allow advanced explorations of the evolution of the biological risk brought by specific stimuli, as cells experience degenerative changes.

**Figure 12:**
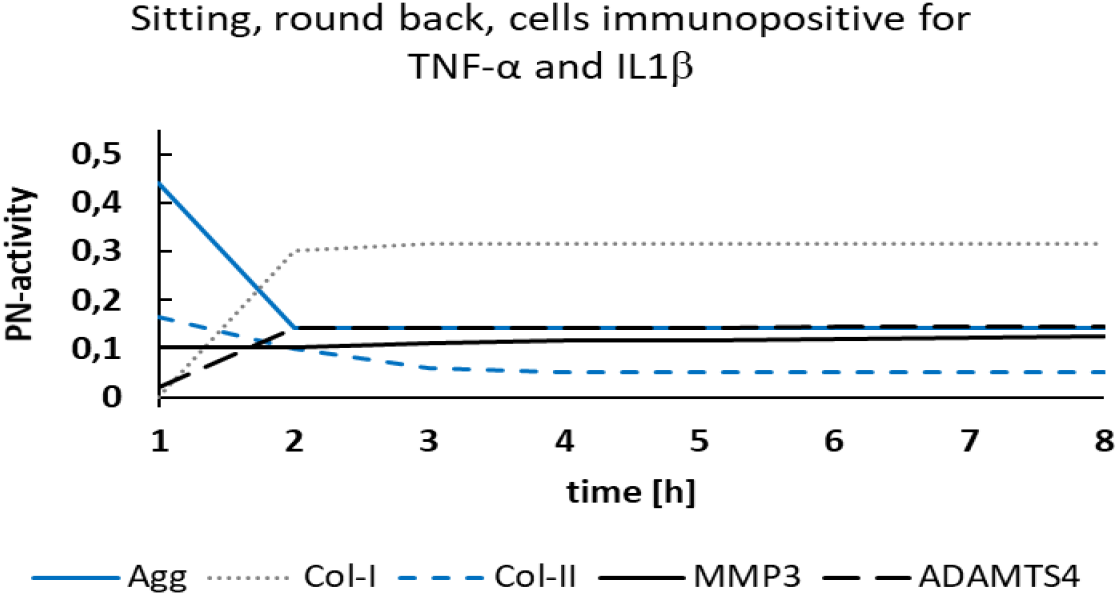
Activity profile of cells exposed to IL1β and TNF-α proinflammatory cytokines during sitting with a round back. Dynamic changes are most obvious in CA for MMP3 mRNA expression, which does not take place in immunonegative cells (Figure 11, B8)

Given the constant evolution of a CA over time, physiologically relevant timeframes must be defined by the user, i.e. the idea of a network model to reach a steady state is intentionally not applicable for time-dependent PN. Arguably, to date, the quality of the results obtained by the PN-Methodology with time-sensitive PN could only be assessed through falsification tests (see Supplementary material 7, 8). This is due to the difficulty to obtain longitudinal experimental data, especially when it comes to non-degenerated IVD tissue. However, the presented results covering several human activities agree well with the knowledge from literature, suggesting that the presented time-sensitivity approach has great potential in network modeling.

## 5. Conclusions

To the best of our knowledge, the PN-Methodology is the first modeling approach that allows a comprehensive approximation of relative cell responses within heterogeneous multifactorial, multicellular systems. Thanks to the interpretation of complex environments as many small feed-forward networks, CA profiles of multiple cell states (or types) can be estimated directly, omitting the simulation of subcellular network interactions. The underlying mathematical approach that interconnects the PN due to equation parameters allows to resolve a system of independent, analytically solvable ODE at low computational cost. Eventually, an option to make PN time-sensitive was presented, allowing dynamic calculations over long periods of time, supplemented by a mathematical solution for transient change of a stimulus nature between activating and inhibiting. The presented network modeling technique with its several underlying methods is deemed to be an important step towards a better understanding of multifactorial disorders that evolve over long periods of time, such as IVD degeneration. Next steps include the application of the PN-Methodology to other cell types and tissues to confirm its potential as a generic theoretical framework to simulate biological systems.

## Acknowledgements

This work was supported by the European Commission and the European Research Council [ERC-2021-CoG-O-Health-101044828], by the Department of Engineering of the Pompeu Fabra University, and by the Spanish Ministry of Science and Innovation and State Agency for Research [STRATO-PID2021-126469OB-C21].

## Author contributions

**Laura Baumgartner:** Conceptualization, Methodology, Software, Validation, Formal analysis, Investigation, Writing-Original Draft, Writing – Review & Editing. **Miguel Ángel González Ballester:** Project administration, Supervision. **Jérôme Noailly:** Resources, Funding acquisition, Project administration, Supervision, Writing – Review & Editing.

## Footnotes

1 Note that each weighting factor is just considered once to determine the size of the system, hence, repetitions of the same weighting factors to determine different CS do not account in the first term of the PN-Equation

## References

1. Stalidzans E, Zanin M, Tieri P, Castiglione F, Polster A, Scheiner S, et al. Mechanistic Modeling and Multiscale Applications for Precision Medicine: Theory and Practice. Netw Syst Med. 2020 May 1;3(1):36–56.

2. Abou-Jaoudé W, Traynard P, Monteiro PT, Saez-Rodriguez J, Helikar T, Thieffry D, et al. Logical modeling and dynamical analysis of cellular networks. Front Genet. 2016 May 31;7(MAY).

3. Panditrao G, Bhowmick R, Meena C, Sarkar RR. Emerging landscape of molecular interaction networks: Opportunities, challenges and prospects. Vol. 47, Journal of Biosciences. Springer; 2022.

4. Baumgartner L, Wuertz-Kozak K, Le Maitre CL, Wignall F, Richardson SM, Hoyland J, et al. Multiscale regulation of the intervertebral disc: Achievements in experimental, in silico, and regenerative research. Int J Mol Sci. 2021;22(2):1– 42.

5. Segarra-Queralt M, Neidlin M, Tio L, Monfort J, Monllau JC, González Ballester MÁ, et al. Regulatory network-based model to simulate the biochemical regulation of chondrocytes in healthy and osteoarthritic environments. Sci Rep [Internet]. 2022;12(3856).

6. Groß A, Kracher B, Kraus JM, Kühlwein SD, Pfister AS, Wiese S, et al. Representing dynamic biological networks with multi-scale probabilistic models. Commun Biol. 2019 Dec 1;2(1).

7. Städter P, Schälte Y, Schmiester L, Hasenauer J, Stapor PL. Benchmarking of numerical integration methods for ODE models of biological systems. Sci Rep. 2021 Dec 1;11(1).

8. Chan SCW, Ferguson SJ, Gantenbein-Ritter B. The effects of dynamic loading on the intervertebral disc. European Spine Journal. 2011;20:1796–812.

9. Segarra-Queralt M, Crump K, Pascuet-Fontanet A, Gantenbein B, Noailly J. The interplay between biochemical mediators and mechanotransduction in chondrocytes: Unravelling the differential responses in primary knee osteoarthritis. Vol. 48, Physics of Life Reviews. Elsevier B.V.; 2024. p. 205–21.

10. Gilbert HTJ, Hodson N, Baird P, Richardson SM, Hoyland JA. Acidic pH promotes intervertebral disc degeneration: Acid-sensing ion channel -3 as a potential therapeutic target. Sci Rep [Internet]. 2016;6(1):1–12.

11. Rinkler C, Heuer F, Pedro MT, Mauer UM, Ignatius A, Neidlinger-Wilke C. Influence of low glucose supply on the regulation of gene expression by nucleus pulposus cells and their responsiveness to mechanical loading. Journal of Neurosurgery: Spine J Neurosurg Spine [Internet]. 2010;13:535–42.

12. Neidlinger-Wilke C, Mietsch A, Rinkler C, Wilke HJ, Ignatius A, Urban J. Interactions of environmental conditions and mechanical loads have influence on matrix turnover by nucleus pulposus cells. Journal of Orthopaedic Research (J Orthop Res). 2012;30(1):112–21.

13. Ju S, Gantenbein-ritter B, Lezuo P, Alini M, Ferguson SJ, Ito K. Effect of Limited Nutrition on In Situ Intervertebral Disc Cells Under Simulated-Physiological Loading. 2009;34(12):1264–71.

14. Illien-Jünger S, Gantenbein-Ritter B, Grad S, Lezuo P, Ferguson SJ, Alini M, et al. The combined effects of limited nutrition and high-frequency loading on intervertebral discs with endplates. Spine (Phila Pa 1976) [Internet]. 2010;35(19):1744–52.

15. MacLean JJ, Lee CR, Alini M, Iatridis JC. Anabolic and catabolic mRNA levels of the intervertebral disc vary … Journal of orthopaedic Research. 2004;22:1193–200.

16. MacLean JJ, Lee CR, Alini M, Iatridis JC. The effects of short-term load duration on anabolic and catabolic gene expression in the rat tail intervertebral disc. Journal of Orthopaedic Research. 2005;23(5):1120–7.

17. Baumgartner L, Sadowska A, Tio L, González Ballester MA, Wuertz-Kozak K, Noailly J. Evidence-based Network Modelling to Simulate Nucleus Pulposus Multicellular Activity in different Nutritional and pro-Inflammatory Environments. Front Bioeng Biotechnol. 2021;9:1–17.

18. Baumgartner L, Reagh JJ, González Ballester MA, Noailly J. Simulating intervertebral disc cell behaviour within 3D multifactorial environments. Bioinformatics. 2021;37(9):1246–53.

19. Kerr GJ, Veras MA, Kim MKM, S?guin CA. Decoding the intervertebral disc: Unravelling the complexities of cell phenotypes and pathways associated with degeneration and mechanotransduction. Semin Cell Dev Biol. 2017;62:94–103.

20. Molinos M, Almeida CR, Caldeira J, Cunha C, Goncalves RM, Barbosa MA. Inflammation in intervertebral disc degeneration and regeneration. J R Soc Interface. 2015;12(104):20141191.

21. Saggese T, Thambyah A, Wade K, McGlashan SR. Differential Response of Bovine Mature Nucleus Pulposus and Notochordal Cells to Hydrostatic Pressure and Glucose Restriction. Cartilage. 2018;00(0):1–13.

22. Johnson ZI, Schoepflin ZR, Choi H, Shapiro IM, Risbud M V. Disc in flames: Roles of TNF-α and IL-1β in intervertebral disc degeneration. Eur Cell Mater. 2015;30:104–17.

23. Le Maitre CL, Freemont AJ, Hoyland JA. The role of interleukin-1 in the pathogenesis of human intervertebral disc degeneration. Arthritis Res Ther. 2005;7(4):R732–45.

24. Purmessur D, Walter B a, Roughley PJ, Laudier DM, Hecht a C, Iatridis J. A role for TNFα in intervertebral disc degeneration: a non-recoverable catabolic shift. Biochem Biophys Res Commun. 2013 Mar;433(1):151–6.

25. Ruiz Wills C, Foata B, González Ballester MÁ, Karppinen J, Noailly J. Theoretical Explorations Generate New Hypotheses About the Role of the Cartilage Endplate in Early Intervertebral Disk Degeneration. Front Physiol [Internet]. 2018;9:1–12.

26. Wilke HJ, Neef P, Caimi M, Hoogland T, Claes LE. New in vivo measurements of pressures in the intervertebral disc in daily life. Spine (Phila Pa 1976) [Internet]. 1999;24(8):755–62.

27. Nachemson A, Elfström G. Intravital dynamic pressure measurements in lumbar discs. Vol. 2, Scandinavian Journal of Rehabilitation and Medication. 1970. p. 1–40.

28. Walsh AJL, Lotz JC. Biological response of the intervertebral disc to dynamic loading. J Biomech. 2004;37(3):329–37.

29. Pachi A, Ji T. Frequency and velocity of people walking. The Structural Engineer. 2005;36–40.

30. Vernillo G, Giandolini M, Edwards WB, Morin JB, Samozino P, Horvais N, et al. Biomechanics and Physiology of Uphill and Downhill Running. Sports Medicine. 2017;47:615–29.

31. Kasra M, Merryman WD, Loveless KN, Goel VK, Martin JD, Buckwalter JA. Frequency Response of Pig Intervertebral Disc Cells Subjected to Dynamic Hydrostatic Pressure. Wiley InterScience. 2006;(October):1967–73.

32. Dudli S, Haschtmann D, Ferguson SJ. Fracture of the vertebral endplates, but not equienergetic impact load, promotes disc degeneration in vitro. Journal of Orthopaedic Research. 2012;30(5):809–16.

33. Gawri R, Rosenzweig DH, Krock E, Ouellet JA, Stone LS, Quinn TM, et al. High mechanical strain of primary intervertebral disc cells promotes secretion of inflammatory factors associated with disc degeneration and pain. Arthritis Res Ther [Internet]. 2014;16(1):R21.

34. Walter B, Korecki C, Purmessur D, Roughley P, Michalek A, Iatridis J. Complex Loading Affects Intervertebral Disc Mechanics and Biology. Osteoarthritis Cartilage. 2012;19(8):1011–8.

35. Mendoza L, Xenarios I. A method for the generation of standardized qualitative dynamical systems of regulatory networks. Theoretical biology & medical modelling (Theor Biol Med Model) [Internet]. 2006;3:13.

36. Belavý DL, Armbrecht G, Felsenberg D. Incomplete recovery of lumbar intervertebral discs 2 years after 60-day bed rest. Spine (Phila Pa 1976). 2011;37(14):1245–51.

37. Nachemson A. Intradiscal measurements of ph in patients with lumbar rhizopathies. Acta Orthopaedica Scandinavica (Acta Orthop Scand). 1969;40(1):23–42.

38. Wilensky U. NetLogo. 1999. p. NetLogo. http://ccl.northwestern.edu/netlogo/.

39. Baumgartner L, Witta S, Noailly J. Parallel networks to predict TIMP and protease cell activity of Nucleus Pulposus cells exposed and not exposed to pro-inflammatory cytokines. bioRxiv 101101/20240828609099 [Internet]. 2024

