## Supplementary material for "Parallel Networks to simulate complex multicellular dynamics - A proof of concept with intervertebral disc cell systems": Baumgartner_et_al_PN_Supplementary_Material.pdf

### Supplementary material 1

#### Calculation of PN-Systems

An example of a complete PN-Equation of the PN-System I of the system of interest of the NP (main article, Figure 4, A) is provided for the Agg mRNA expression of cells immunopositive for IL1β, for an initially anabolic mag and catabolic freq (Figure S1.1).

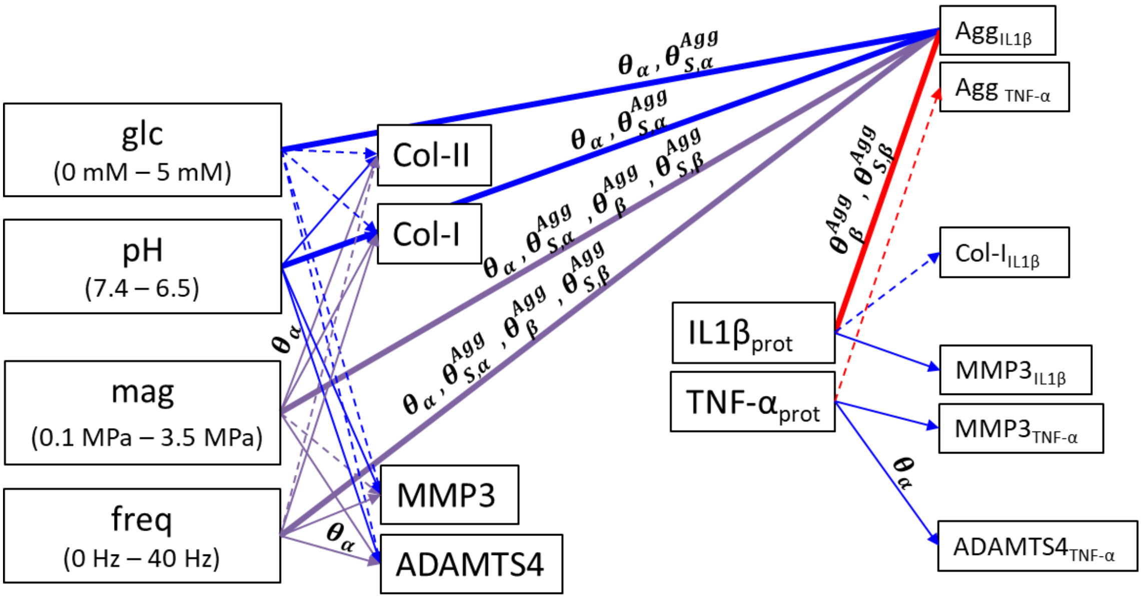


$\theta_\alpha$: all (potentially) activating weighting factors within the PN-System

$\theta_{S,\alpha}^{Agg}$: activating weighting factors that are directly acting on Agg of IL1β inflamed cells

$\theta_{\beta}^{Agg}$: all (potentially) inhibiting weighting factors of Agg, independently of the CS

$\theta_{S,\beta}^{Agg}$: inhibiting weighting factors that are directly acting on Agg of IL1β inflamed cells

$$\frac{d\omega_{Agg,\,IL1\beta\,infl}}{dt} = \left(\left(\frac{1+\sum\theta_\alpha}{\sum\theta_\alpha}\right)\left(\frac{\theta_{glc,\alpha}^{Agg}x_{glc,\alpha}^{Agg}+\theta_{pH,\alpha}^{Agg}x_{pH,\alpha}^{Agg}+\theta_{mag,\alpha}^{Agg}x_{mag,\alpha}^{Agg}}{1+\theta_{glc,\alpha}^{Agg}x_{glc,\alpha}^{Agg}+\theta_{pH,\alpha}^{Agg}x_{pH,\alpha}^{Agg}+\theta_{mag,\alpha}^{Agg}x_{mag,\alpha}^{Agg}}\right)\right)\cdot$$
$$1-\left(\left(\frac{\theta_{\delta,\,mag,\beta}^{Agg}+\theta_{freq,\beta}^{Agg}+\theta_{IL1\beta,\beta}^{Agg}}{(\boldsymbol{\theta_{glc,\alpha}^{Agg}}+\boldsymbol{\theta_{pH,\alpha}^{Agg}})+(\boldsymbol{\theta_{\delta,mag,\beta}^{Agg}}+\boldsymbol{\theta_{IL1\beta,\beta}^{Agg}}+\boldsymbol{\theta_{freq,\beta}^{Agg}})}\right)\cdot\left(\frac{1+\theta_{\delta,\,mag,\beta}^{Agg}+\theta_{freq,\beta}^{Agg}+\theta_{IL1\beta,\beta}^{Agg}+\theta_{TNF-\alpha,\beta}^{Agg}}{\theta_{\delta,\,mag,\beta}^{Agg}+\theta_{freq,\beta}^{Agg}+\theta_{IL1\beta,\beta}^{Agg}+\theta_{TNF-\alpha,\beta}^{Agg}}\right)\right.$$
$$\left.\cdot\left(\frac{\theta_{\delta,\,mag,\beta}^{Agg}x_{mag,\beta}^{Agg}+\theta_{freq,\beta}^{Agg}x_{freq,\beta}^{Agg}+\theta_{IL1\beta,\beta}^{Agg}x_{IL1\beta,\beta}^{Agg}}{1+\theta_{\delta,\,mag,\beta}^{Agg}x_{mag,\beta}^{Agg}+\theta_{freq,\beta}^{Agg}x_{freq,\beta}^{Agg}+\theta_{IL1\beta,\beta}^{Agg}x_{IL1\beta,\beta}^{Agg}}\right)\right) \tag{S1.1}$$

$$with$$

$$\sum\theta_\alpha = \theta_{glc}^{Agg}+\theta_{pH}^{Agg}+\theta_{glc}^{ColII}+\theta_{pH}^{ColII}+\theta_{glc}^{ColI}+\theta_{pH}^{ColI}+\theta_{glc}^{MMP3}+\theta_{pH}^{MMP3}+\theta_{glc}^{ADAMTS4}+\theta_{pH}^{ADAMTS4}+\theta_{IL1\beta}^{ColI}+\theta_{IL1\beta}^{MMP3}$$
$$+\theta_{TNF-\alpha}^{MMP3}+\theta_{TNF-\alpha}^{ADAMTS4}+\theta_{mag}^{Agg}+\theta_{freq}^{Agg}+\theta_{mag}^{ColII}+\theta_{freq}^{ColII}+\theta_{mag}^{ColI}+\theta_{freq}^{ColI}+\theta_{mag}^{MMP3}+\theta_{freq}^{MMP3}+\theta_{mag}^{ADAMTS4}+\theta_{freq}^{ADAMTS4}$$

Figure S1.1: Above: PN-Equation for the Agg mRNA expression of IL1β inflamed cells. Every possible contribution of the weighting factors is illustrated. $\boldsymbol{\theta_\alpha}$ is illustratively represented in the PN-System but stands for all potentially activating S-CA relationships, see $\sum\boldsymbol{\theta_\alpha}$ of the equation S1. Below: Eq. S1.1 reflecting the Agg mRNA expression under an initially anabolic mag and catabolic freq. $\boldsymbol{\delta}$ reflects completive weighting factors.

Note that time-dependent weighting factors must be mentioned only once in the denominator of the first inhibiting term (Figure S1.1, bold), to accurately reflect the relative weight of inhibition with respect to activation.

**PN-Systems associated with the local stimuli IL1β and TNF-α (PN-Systems II & III, in main article, Figure 4, A)**

PN-Systems II and III only contain one parallel network each. Therefore, the first fraction of the activating part of their respective PN-Equations only considers the weighting factors related to the specific CA (i.e. the CA for the mRNA expressions of the local stimuli IL1β and TNF-α). Eq. S1.2 shows the PN-Equation for TNF-α:

$$\frac{d\omega_{TNF-\alpha,CS}}{dt} = \left(\left(\frac{1+\theta^{TNF-\alpha}_{glc,\alpha}+\theta^{TNF-\alpha}_{pH,\alpha}+\theta^{TNF-\alpha}_{(\delta),mag,\alpha}+\theta^{TNF-\alpha}_{(\delta),freq,\alpha}}{\theta^{TNF-\alpha}_{glc,\alpha}+\theta^{TNF-\alpha}_{pH,\alpha}+\theta^{TNF-\alpha}_{(\delta),mag,\alpha}+\theta^{TNF-\alpha}_{(\delta),freq,\alpha}}\right)\left(\frac{\sum\theta^{CA}_{S,\alpha}x^{CA}_{S,\alpha}}{1+\sum\theta^{CA}_{S,\alpha}x^{CA}_{S,\alpha}}\right)\right) \cdot \left(1-\left(\left(\frac{\sum\theta^{CA}_{S,\beta}}{\sum\theta^{CA}_{S,\alpha}+\sum\theta^{CA}_{S,\beta}}\right)\left(\left(\frac{1+\sum\theta^{CA}_{\beta}}{\sum\theta^{CA}_{\beta}}\right)\left(\frac{\sum\theta^{CA}_{S,\beta}x^{CA}_{S,\beta}}{1+\sum\theta^{CA}_{S,\beta}x^{CA}_{S,\beta}}\right)\right)\right)\right) \qquad \text{(S1.2)}$$

#### Supplementary material 2

#### Overview of weighting factors considered in the system of interest of the NP

The subsequent Table S2.1 contains an overview of the weighting factors used to calculate the current results. The information for indirect mechanotransduction was previously presented, together with the mathematical description of weighting factors (1). In short: weighting factors $\theta_S^{CA}$ were obtained from experimentally found x-fold mRNA expressions ($\epsilon$). $\epsilon$ were translated into a "cellular effort" ($f(\epsilon)$), which allows to equalize up- and downregulations of mRNA expressions compared to a control value. $f(\epsilon)$, of significant changes in x-fold mRNA expressions were then normalized to obtain the weighting factor $\theta_S^{CA}$. Non-significant x-fold mRNA expressions were assigned to a weighting factor of $\theta_S^{CA}$ = 0.0100.

Table S2.1: Overview of weighting factors for the tackled system of interest of the NP. Individual weighting factors were derived from the cellular effort ($\boldsymbol{f(\epsilon)}$), based on x-fold mRNA expressions ($\boldsymbol{\epsilon}$). The scaling factor $\boldsymbol{\vartheta_{\theta_{max}}}$= $\boldsymbol{\vartheta_1 = 28.7}$ was determined by the S-CA relationship pH-MMP3. NS: Not significant; act: activating; inh: inhibiting; *: estimated $\boldsymbol{\epsilon}$; glc: glucose.

| **Stimulus** | **mRNA** | $\boldsymbol{\epsilon}$ | $\boldsymbol{f(\epsilon)}$ | $\boldsymbol{\theta_S^{CA}}$( $\vartheta_1$ = 28.7) | **Source** | **Cell type** |
|---|---|---|---|---|---|---|
| glc | Agg | NS, act | - | 0.0100 | (2) | human |
| | Col-I | NS, act | - | 0.0100 | (2) | human |
| | Col-II | NS, act | - | 0.0100 | (2) | human |
| | MMP3 | NS, act | - | 0.0100 | (2) | human |
| | ADAMTS4 | NS, act | - | 0.0100 | (1) | bovine |
| | IL1β | NS, act | - | 0.0100 | (1) | bovine |
| | TNF-α | NS, act | - | 0.0100 | (1) | bovine |
| pH | Agg | 0.37 | 2.7027 | 0.0942 | (3) | human |
| | Col-I | NS, act | - | 0.0100 | (3) | human |
| | Col-II | 0.63 | 1.5873 | 0.0553 | (4) | bovine |
| | MMP3 | 28.7 | 28.7000 | 1.0000 | (3) | human |
| | ADAMTS4 | 5.7 | 5.7000 | 0.1986 | (3) | human |
| | IL1β | 81 | 81.0000 | 2.8223 | (3) | human |
| | TNF-α | NS, act | - | 0.0100 | (3) | human |
| IL1β | Agg | 0.45* | 2.2222 | 0.0774 | (5) | human |
| | Col-I | NS, act | - | 0.0100 | (5) | human |
| | Col-II | NS, inh | - | 0.0100 | (5) | human |
| | MMP3 | 10.8* | 10.8000 | 0.3763 | (5) | human |
| | ADAMTS4 | NS, inh | - | 0.0100 | (5) | human |
| TNF-α | Agg | NS, inh | - | 0.0100 | (1) | bovine |

| | | | | | | |
|---|---|---|---|---|---|---|
| | Col-I | 0.31 | 3.2258 | 0.1124 | (1) | bovine |
| | Col-II | 0.06 | 16.6667 | 0.5807 | (1) | bovine |
| | MMP3 | 26.85 | 26.8500 | 0.9355 | (1) | bovine |
| | ADAMTS4 | 5.77 | 5.7700 | 0.2010 | (1) | bovine |
| Mag | Agg | 10 | 10 | 0.3484 | (6) | rat |
| | Col-I | 25 | 25 | 0.8711 | (6) | rat |
| | Col-II | 4 | 4 | 0.1394 | (6) | rat |
| | MMP3 | NS | - | 0.0100 | (6) | rat |
| | ADAMTS4 | 4 | 4 | 0.1394 | (6) | rat |
| | TNF-α | NS | - | 0.0100 | Indications (7,8) | |
| | IL1β | NS | - | 0.0100 | Indications (9) | |
| Freq | Agg | 10<br>1.05* | $\frac{10}{1.05}$<br>(= 9.5238) | 0.3318 | (6) | rat |
| | Col-I | 25<br>0.50* | $\frac{25}{2}$<br>(=12.5000) | 0.4355 | (6) | rat |
| | Col-II | NS | - | 0.0100 | (6) | rat |
| | MMP3 | 15<br>1.02* | $\frac{15}{1.02}$<br>(=14.7059) | 0.5124 | (6) | rat |
| | ADAMTS4 | 8<br>2.66 | $\frac{8}{2.66}$<br>(=3.0075) | 0.1048 | (6) | rat |
| | TNF-α | NS | - | 0.0100 | No information found | |
| | IL1β | NS | - | 0.0100 | No information found | |

### Supplementary material 3

#### Development of the PN-Equation

The PN-Equation is developed based on a well-established graph-based method presented by Mendoza and Xenarios (10) (Eq. S3.1 & S3.2).

$$\frac{dx_i}{dt} = \frac{-e^{0.5h} + e^{-h(\omega_i - 0.5)}}{(1 - e^{0.5h})(1 + e^{-h(\omega_i - 0.5)})} - \gamma_i x_i \qquad \text{(S3.1)}$$

$$with$$

$$\omega_i = \left(\frac{1 + \sum \alpha_n}{\sum \alpha_n}\right)\left(\frac{\sum \alpha_n x_n^a}{1 + \sum \alpha_n x_n^a}\right)(1 - \left(\frac{1 + \sum \beta_m}{\sum \beta_m}\right)\left(\frac{\sum \beta_m x_m^i}{1 + \sum \beta_m x_m^i}\right)) \qquad \text{(S3.2)}$$

Eq. S3.1 determines the overall activation of a node, $x_i$, where the gain variable ($h$) allows to gradually shape the relationship between $x_i$ and the total nodal input, $\omega_i$, from a linear to a step function. The overall node activation can be reduced by integrating a decay rate (γ). Eq. S3.2 expresses the total input of the node, according to the set of activators $x_n^a$ and inhibitors $x_m^i$ that act on the node with the respective positive strengths $\alpha_n$ and $\beta_m$ (Figure S3.1, A). Nodal activation is always bound between 0 and 1.

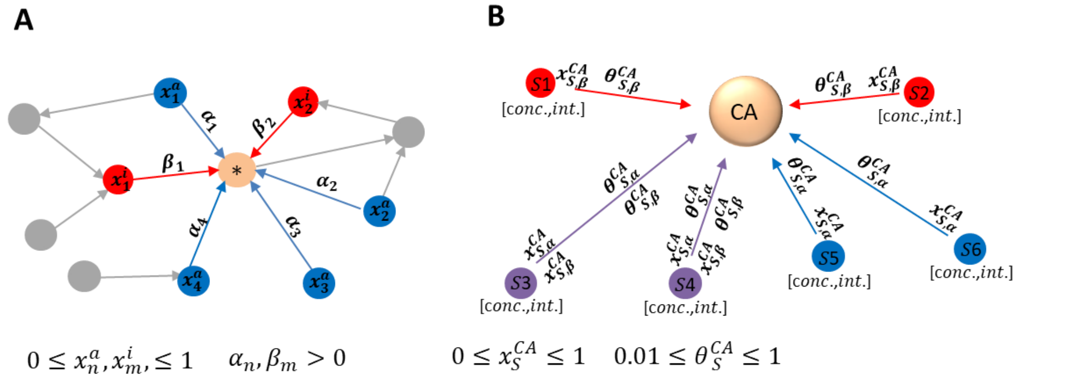


Figure S3.1: Comparison between the network presented by Mendoza and Xenarios 2006 (10) (A) and the concept of parallel network modelling to tackle heterogeneous stimulus environments (B), where an individual parallel network for a given stimulus environment of six stimuli is shown. The overall activity of the principal node (* vs. CA) is determined by adjacent network nodes (A), or by a local stimulus environment (B). The effect of adjacent nodes is either activating (blue, α) or inhibiting (red, β) (A, B) or includes dose and time dependencies (B, purple; S3, S4). Whilst network nodes are usually interconnected within each other (A), the objective of individual parallel networks (B) is to determine a final CA. $\boldsymbol{\alpha_n, \beta_m, \theta_S^{CA}}$: activating/inhibiting factors. $\boldsymbol{x_n^a, x_m^i}$: adjacent node activation (A), $\boldsymbol{x_S^{CA}}$: sensitivity to a stimulus dose, i.e. concentration (conc.) /intensity (int.) (B).

A PN-Network is always directional as it reflects the effect of adjacent nodes on one principal node that stands for a CA (Figure S3.1, B). Adjacent nodes represent the stimulus (micro-) environment. The impact of each adjacent node is determined by the product of a weighting factor ($\theta_S^{CA}$) and the corresponding stimulus concentration/intensity ($x_S^{CA}$). Moreover, the integration of time- and dose-dependencies within the PN-Network allows a stimulus to be either activating or inhibiting, according to the user-defined stimulus dose and the time, during which the condition is maintained (see main article).

Within the PN-Methodology, the decay term defined by Mendoza & Xenarios (Eq. S3.1) would reflect a possible decay in mRNA expression under sustained stimulation or the half-life of a secreted protein. In the present methodology the decay terms would represent, however, an additional source of uncertainty, and they don’t add further information about the regulation of the system of interest (which tackles CA in terms of mRNA expression or protein synthesis, rather than actual amounts of mRNA or proteins in the system) and were, therefore, not considered. Moreover, given that CA are directly obtained through the S-CA relationships that are based on experimental findings, no additional manipulation of $\omega_i$ is needed, i.e. $h$ in Eq. S3.1 reflects a linear relationship between $x_i$ and the total nodal input $\omega_i$ in the PN-Methodology (please check original research with regard to linearity (10)).

Accordingly, the overall CA was directly assessed by $\omega_i$, leading to Eq. S3.3, which reflects the basis for the PN-Equation.

$$\frac{dx_i}{dt} = \frac{d\omega_i}{dt} \qquad \text{(S3.3)}$$

### Supplementary material 4

#### Estimation of the adjustment factor ($\sigma$) to approximate mRNA expressions in NP cells

To estimate $\sigma$, it was supposed that 8h of axial dynamic loading of 0.2-0.8 MPa and 0.1-1 Hz may not induce degeneration of the disc (11). Hence, model predictions for 0.2 MPa and 0.8 MPa at a medium freq of 0.5 Hz were calculated for adjustment factors of $\sigma = 1$ to $\sigma = 4$ (non-inflamed cells, under optimal, nutritional conditions of 5 mM glc, pH 7.1) and results were evaluated with regard to their catabolic progression at 0.8 MPa (Figure S4.1).

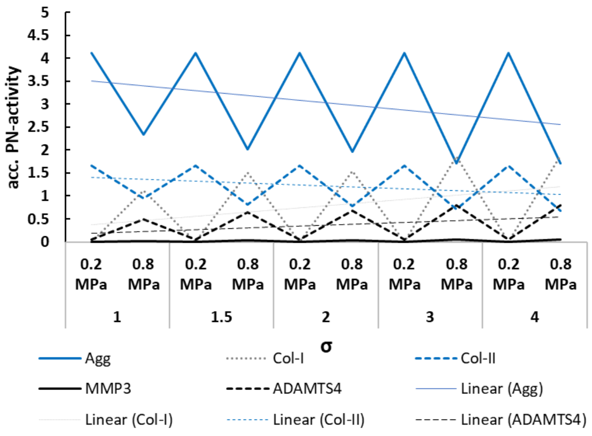


Figure S4.1: model predictions for different adjustment factors ($\boldsymbol{\sigma}$) and corresponding tendencies. Optimal nutritional conditions (5 mM glc, pH 7.1) and a frequency of 0.5 Hz were used.

Results showed that catabolic predictions at 0.8 MPa were already elevated for $\sigma = 2$. This is reflected by a higher acc. PN-activity for Col-I than Col-II, a considerably lower Agg mRNA expression compared to $\sigma = 1$ and a close approximation of the CA of Col-II and ADAMTS4. Hence, as an initial approximation for an adjustment factor, a medium value between $\sigma = 1$ and $\sigma = 2$, i.e. $\sigma = 1.5$ was set. This adjustment factor was implemented for both, mag and freq, but shall be individually determined as soon as adequate experimental information is available.

#### Supplementary material 5

**Estimation of individual time sensitivities ($\gamma_S^{CA}$) of mRNA expressions in NP cells**

$\gamma_S^{CA}$ within the anabolic range is referred to as $\gamma_{S,ana}^{CA}$ and within the catabolic range as $\gamma_{S,cata}^{CA}$. $\gamma_{S,i,ana}^{CA}$ was estimated based on experimental data, where the effect of 1 MPa and 1 Hz on rat NP cells was assessed after 0.5h, 2h and 4h (12). Thereby, experimental data was translated into a mathematically assessed "cellular effort" as done for the determination of the weighting factors (previously explained (1)) and the most pronounced change in CA due to the effect of time was considered. $\gamma_{S,i,cata}^{CA}$ was derived from the values assessed for $\gamma_{S,i,ana}^{CA}$. Thereby, Agg was chosen as a reference value (1) to determine the individual time delays of CA to unfold their whole catabolic potential. Time sensitivities within the catabolic range for any other CA were obtained as fractions, relative to Agg (Table S5.1).

Table S5.1: individual time sensitivities for initially anabolic (ana) and catabolic (cata) stimulus intensities. Agg was used as a reference for the determination of catabolic time sensitivities.

| **CA** | $\gamma_{mag,i,ana}^{CA}/\gamma_{freq,i,ana}^{CA}$ | $\gamma_{mag,i,cata}^{CA}/\gamma_{freq,i,cata}^{CA}$ |
|---|---|---|
| Agg (ref) | 3.1429 | 1 |
| Col-II | 1.7500 | $\frac{1.7500}{3.1429} = 0.5568$ |
| Col-I | 2.7647 | $\frac{2.7647}{3.1429} = 0.8797$ |
| MMP3 | 2.1571 | $\frac{2.1571}{3.1429} = 0.6864$ |
| ADAMTS4 | 3.1481 | $\frac{3.1481}{3.1429} = 1.0017$ |
| TNF-α | 0.3143 | No catabolic time sensitivity |
| IL1β | 0.3143 | No catabolic time sensitivity |

With regard to proinflammatory cytokines, no information about a possible time-sensitivity to loading conditions was found. Therefore, prudent time sensitivities were initially chosen for $\gamma_{S,i,ana}^{CA}$, being 10 times lower than the time sensitivity of Agg. This assumption was based on the observation that long-term exposure to microgravity does not categorically lead to IVD failure (13), whereas excessive inflammatory conditions might provoke non-recoverable catabolic shifts (14). Under initially catabolic loading, no time sensitivity was programmed, thus, fixed values directly provided by the generic function were used to determine the proinflammatory environment.

### Supplementary material 6

**Qualitative validation of the PN-Methodology: simulation of exposure to microgravity and physical activities**

The duration of exposure to microgravity was derived from expeditions to the International Space Station (ISS) launched by the NASA (15). The effect of exposure to microgravity was compared to the effect calculated through an approximation of a corresponding time spent on Earth. An average day was thereby distributed in 10h sleeping/laying down (night/day rest), two episodes of 4h sedentary work (sitting) and three episodes of 2h physical activity (walking). Hence, the simulated day reflects an optimal human moving behavior. The accumulated mRNA expression over one entire day was calculated by adding the mRNA expressions of each physical activity. Thereby, the period of time, during which a physical activity was carried out, matters but not its sequence, e.g. the CA for Agg mRNA expression of one day is the sum of accumulated CA for Agg mRNA expression after 10h resting/sleeping, plus two times the accumulated CA for Agg mRNA expression after 4h sitting, plus three times the accumulated CA for Agg mRNA expression after 2h walking. Given the calculation time step of 1h, the CA was estimated after each hour of a certain physical activity. Due to the constant boundary conditions that determine physical activity of one day, the accumulated mRNA expression over 180 days was obtained by multiplying the accumulated mRNA expression of one day by 180. Simulations for four different CS (non-inflamed and inflamed for TNF-α, IL1β and both, TNF-α&IL1β) were provided (see Results, main article, and supplementary material 8).

Intradiscal pressures for both, exposure to microgravity and physical activities were obtained from *in vivo* measurements (16) and rounded to the minimal step size of loading parameters of 0.05. Whenever intradiscal pressures were provided as ranges, average values were used for the simulation (e.g. walking: 0.53 MPa – 0.65 MPa (16) was considered as 0.60 MPa). For microgravity calculations, the intradiscal pressure reflected the (rounded) value after 7h of rest (16). Freq for hiking was estimated to be around 1 Hz, whilst values for walking and jogging were within the range as found in observational studies (17,18). A vibration for “sitting in/on a motor vehicle” was estimated within the range of maximal catabolic predictions. An overview of the simulated human activities with corresponding loading parameters and duration is provided in the main article (Table 4).

### Supplementary material 7

#### 7.1 The IVD case study - case-specific implementations

The PN-Methodology was developed to investigate initiations of Intervertebral disc (IVD) degeneration. Thereby, the presented results aim to contribute to a better understanding of the activity of Nucleus Pulposus (NP) cells by modeling dose- and time-dependent cell regulatory processes that are extremely difficult to measure directly either *in vitro* or *in vivo*.

To investigate initiations of IVD degeneration, we opted for a comparison of CA in terms of mRNA expressions of ECM proteins and important proteases under different combinations of key relevant, external stimuli and proinflammatory cytokines. As a consequence, the effective degradation of ECM proteins due to MMP3 and ADAMTS4 enzymes is not considered. Yet, this case study allows for qualitative interpretations of catabolic activities in the IVD, where increased protease mRNA expression would deplete the ECM, on top of the downregulation of ECM protein mRNA expression. If a quantification of the additional effect of synthesized proteases on ECM would be required, it could be obtained by creating a new system of interest; roughly, by considering MMP3 and ADAMTS4 as local stimuli (i.e., instead of TNF-α and IL1β; see main article, Figure 3, below), and feeding the system with information about protein synthesis, rather than mRNA expression.

The programming of time-related dynamics for the IVD consider a loss of anabolic stimulus over time and a time delay in the development of a catabolic responses to a catabolic stimulus. The latter was programmed to take into account that short durations of high loading impacts might not promote IVD degeneration (7), whilst sustained overload as such is considered as a risk factor for degenerative changes (19).

#### 7.2 Relevance of the obtained results for the NP tissue

**Predictions of daily moving habits (main article, Figure 11)**

Fluctuating mechanical stimuli affect the CA in terms of both tissue proteins and protease mRNA expression. Thereby, the cell response to different combinations of stimulus concentrations does not seem to follow a systematic pattern per-se, especially for the motions expected to involve catabolic cell activities (CA). For example, while two-hours sitting with a round back and hiking with extra weight were predicted to particularly upregulate the mRNA expressions of Collagen Type I (Col-I), followed by ADAMTS4, jogging and vibration might rather trigger MMP3 mRNA expressions (main article, Figure 11, B1).

Physical activities that are reported to be less related with IVD degeneration were sitting (office workers) (20) and walking (21,22). In contrast, activities classically related to IVD degeneration or to catabolic changes in NP cell responses were weightlifting (23), heavy work (carpenters) (20) or exposure to whole-body vibration (machine drivers) (20). Model simulations are in agreement with such findings, as they predict walking and hiking without extra weight to be highly anabolic, followed by sleeping and sitting with active back (main article, Figure 11, B1, B2-B5). Accordingly, walking with extra weight or vibrations were predicted by the model to cause catabolic cell responses (main article, Figure 11, B1, B6, B9). However, hiking does not seem to be specifically linked to IVD degeneration, which could be explained with the use of supporting hip belts for (heavy) backpacks. Hence, hikers most probably carry heavy weights rather on their hips than on their shoulders and CA profiles of hikers generally rather resemble the predictions for "hiking without extra weight" (main article, Figure 11, B1, B5). Interestingly, sitting with a round back was also catabolic in contrast to sitting with an active back (main article, Figure 11, B3 vs. B8), which nicely corresponds to current ergonomic paradigms.

A limitation related with the modelling of the effect of magnitude (mag) and frequency (freq) is that rodent experimental data had to be used to estimate the weighting factors and time dependencies of different CA. Moreover, there is also a lack of further individualization of time-sensitivities and adjustment factors between mag and freq, due to the limited availability of experimental data for quantification. Likewise, more experimental evidence about the relationship between proinflammatory cytokines and (physiological) loading parameters would contribute to confirm or adapt current modelling assumptions. Eventually, to determine weighting factors, ideally, the maximal effect of a stimulus within a physiological range on a CA should be addressed in experimental studies to be fully exploited in the model. For example, the available data about the mechanostimulation were measured at 1 MPa and 0.01 Hz, 0.2 Hz and 1 Hz, respectively (6). *In vivo* measurements in humans clearly indicate that the physiological range is much broader (16,24). Consequently, our weighting factor for mag and freq might have been underestimated compared to the effect of nutrient environments, for which measurements over the whole physiological range were available.

Whilst IVD aging appears to be a physiological adaptation of the tissue, accelerated IVD degeneration is not affecting everyone, despite the catabolic shifts possibly caused by moving habits that many people experience more or less frequently (e.g. carrying heavy weights on the shoulders, exposure to vibration). Assuming that acute physiological moving habits are well regulated by the human organism, stimulus chronicity and, therefore, the factor time, seems to be important in IVD degeneration. Furthermore, the high diversity of cellular responses due to different combinations of stimuli suggests that the mechanisms finally leading to (micro-) injuries are manifold, without following one general systematic pattern, which would go along with the high variability of phenotypes in IVD degeneration.

The impact of chronicity is particularly interesting in the model predictions for jogging (main article, Figure 11, B7). The anabolism of jogging has a strongly time dependent component.

Whilst its accumulated Parallel Network (PN)-activity was predicted to be considerably lower than e.g. walking (main article, Figure 11, B1), insights into the evolution over time (main article Figure 11, B7) suggest that the catabolic impact is pronounced after two hours of jogging, whilst jogging for one hour is still largely anabolic. Accordingly, findings in literature regarding the effect of jogging are controversial: whereas a study of young adults found relationship between frequent jogging and degenerative changes in lumbar IVD (21), other findings did not reveal significant effects (25) or jogging was found to be beneficial compared to a non-physiologically active control group (22,26). Assuming that the chronicity of cell stressors is crucial in the dynamics of IVD degeneration, as suggested by the model predictions, a frequent switch among different moving habits can contribute to reduce the risk of IVD degeneration and/or decelerate its development.

**Predictions of the effect of microgravity and daily life under gravity and for different proinflammatory cell states (CS)**

Simulations of a daily life on Earth show low to minimal Col-I and protease mRNA expressions and elevated Aggrecan (Agg) and Collagen Type II (Col-II) mRNA expressions (Figure S7.1, A2).

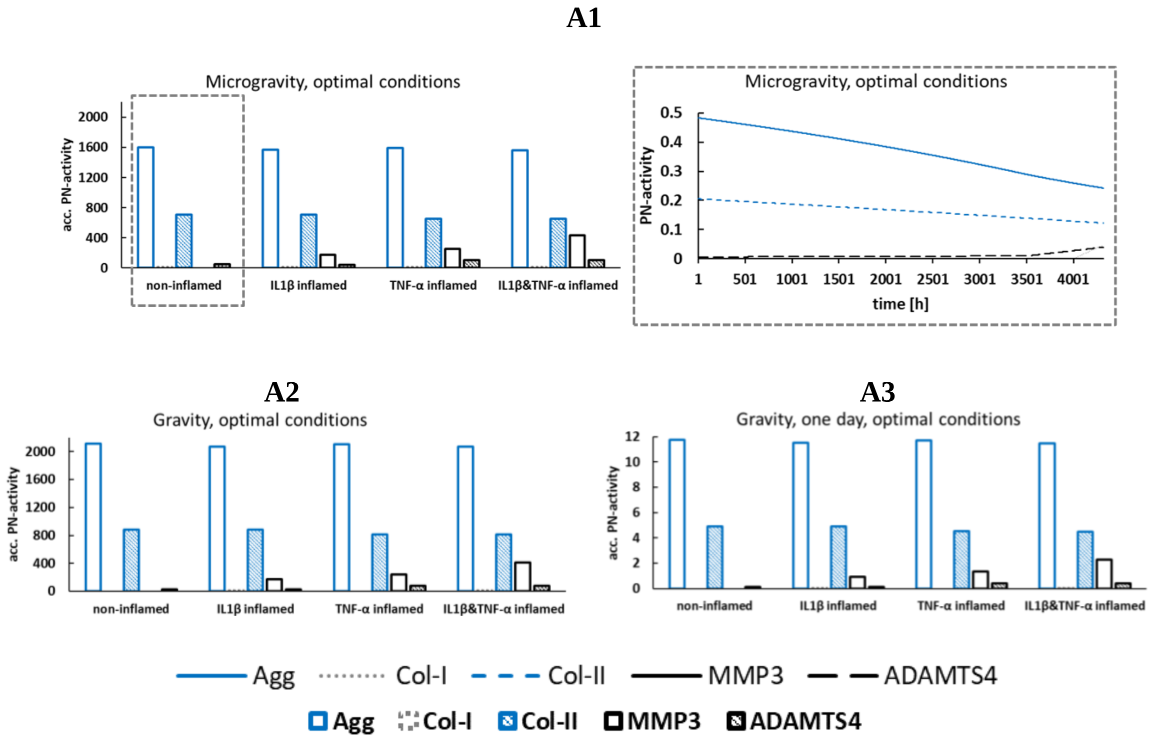


Figure S7.1: estimated CA profiles after six months of exposure to microgravity (A1, left), for an average daily moving habit (A3) and its accumulation over six months (A2). The effect of exposure to microgravity is additionally shown for cells that are not influenced by proinflammatory cytokines as a continuous development over time (A1, right). Non-degenerated nutritional conditions were used (5 mM glucose, pH 7.1). Please note that the graphs A1 and A2 were provided in the main article, Figure 10, and were hereby repeated for the context of graph A3.

The presence of proinflammatory cytokines causes protease mRNA expressions to rise, whilst the mRNA expressions of tissue proteins show an overall slight decrease, independently of the simulated condition. Thereby, the predicted catabolic shift in TNF-α immunopositive cells is in agreement with experimental findings that identify TNF-α as a strong catabolic mediator (14). IL1β was predicted to be less catabolic than TNF-α, which coincides with its possible role in normal cell homeostasis (27). Also independently of the simulated condition, Col-II mRNA expression is lower, relative to Agg mRNA expression (Figure S7.1). Lower CA for Col-II than for Agg along with both, a lower tissue turnover of collagen compared to Agg (28,29) and a lower fraction of Col-II tissue compared to Agg (30).

Model predictions simulating daily life under gravity (Figure S7.1, A2) were in agreement with general expectations: minimal or close to minimal CA for MMP3 and ADAMTS4 mRNA expressions agree with low MMP3 and ADAMTS4 protease levels found within the tissue (31). Elevated CA for Agg and Col-II mRNA expressions reflect the cellular function to maintain the tissue.

Exposure to microgravity over six months led to a general downregulation of the expression of tissue proteins by predicting a roughly 25% and 20% lower PN-activity of Agg and Col-II mRNA expression, respectively, independently of the CS (Figure S7.1, A1, left vs. A2). Thereby, a downregulation of glycosaminoglycan contents is supported by various animal models (13). Agg is the major proteoglycan in cartilaginous tissues. It is a complex macromolecule with numerous chondroitin sulfate glycosaminoglycan chains. Because these chains are negatively charged, Agg is responsible to attract water through Donnan osmosis effects. A reduced Agg content goes along with a reduced fixed charge density, which in turn could be related to a lower hydration of the IVD (30). Although such findings at first glance would explain disc desiccations found after long-term space flights (32), the slow turnover of tissue proteins with half-lives of around 12 (Agg) and 95 (collagen) years (28,29), questions whether the effect of tissue protein downregulation becomes visible *in vivo* at the tissue level, after only six months.

MMP3 mRNA expression was not particularly affected by simulations of microgravity. In contrast, ADAMTS4 mRNA expression was predicted to slightly rise, which could be generally attributed to the end of the stay in space, i.e. after around 3500 h (Figure S7.1, A1, right). With this regard, the value of the adjustment factor is deterministic for the velocity of the continuous evolution of CA profiles over time. Given that the adjustment factor could not be specified yet between mag and freq, and that rather the range, than its effective size was determined (Supplementary Material 4), a generally prudent interpretation of the specific timepoints observed in continuous CA-profiles is required.

Remarkably, though, the continuous evolution of a CA over time (Figure S7.1, A1, right) shows a general trend of a constant low level of proteases within the first months of simulated exposure to microgravity. Such expressions were similar to one simulated day under gravity (Figure S7.1, A3). The daily moving habit was chosen to be quite "optimal", while "suboptimal" moving under gravity (i.e. carrying heavy weights or exposure to vibration) was responsible for the upregulation of protease activity (main article, Figure 11, B6-B9). Under microgravity such "suboptimal" stimuli cannot be represented; consequently, the protease mRNA expression was kept low, which might contribute to an increased swelling behavior of the IVD along time. Swelling behavior of the IVD was particularly observed under head-down-tilt bedrest (33), the most common analogue to investigate effects of microgravity. Yet, the existence of IVD swelling during spaceflight is debated, since studies found that pre-to post-flight differences in disc water content after six months exposure to microgravity were either non-significant (34) or even signs for disc desiccation were observed already during exposure to microgravity in ultrasound images (32).

Hence, the model predicts indications for both, disc desiccation on the one hand, due to a downregulation of tissue protein mRNA expression, whilst, on the other hand, constantly low levels of protease mRNA expression might contribute to findings of disc swelling. Both is actually observed under similar mechanical conditions, i.e. under exposure to microgravity and in bed rest studies. A better understanding of a final consolidation at the tissue level might be obtained by a quantification of actual mRNA expressions coupled to a CA. Moreover, besides the mechanical loading conditions, peculiarities of microgravitational conditions, e.g. lumbar spine flattening and possible changes in intradiscal nutrient supply should be considered in future work to confirm or adapt current predictions under microgravity.

#### Conclusions of the tissue-specific findings

This work is the first contribution that allows to describe relative cell responses within complex, heterogeneous stimulus environments. The aim was to approximate cell responses to a native stimulus environment by considering crucial external and local stimuli. Thanks to the integration of time-dependent effects, this work uniquely allowed to estimate adaptations of cells to persisting mechanical load patterns over time, as found in daily activities or under exposure to microgravity.

Findings were in good agreement with literature and coincide with observed organ and tissue macroscopic adaptations. Hence, simulations with the PN-Methodology indicate that IVD NP degeneration is most probably not the result of one, principal biological response, but it highly depends on the (micro-) environmental stimulus combinations. Results further suggest that chronicity of mechanical loads is key relevant in IVD NP degeneration. Hence, a frequent change of moving habits, e.g. sitting positions, might be beneficial to maintain IVD integrity.

Future work should tackle further detail regarding the effects of exposure to microgravity and it should focus on a more sensitive integration of loading conditions. Eventually, an integration of this approach into (bottom-up or top-down) multiscale approaches is deemed to be key relevant to tackle injury dynamics within the IVD.

#### Supplementary material 8

##### Additional simulations: modelling nutritional conditions as found in early degenerated IVD and predictions of the PN-Methodology coupled to a 3D Agent-based model, simulating a multicellular environment

Methods: The most critical conditions expected in early degenerated IVD were simulated. Therefore, the region of the NP was chosen where the most adverse nutrient environment was found. Its location and local nutrient concentrations were determined by our in-house mechanotransport finite element (FE) model (35); the cartilage endplate permeability was adapted as expected at early stages of IVD degeneration and nutrient concentrations were estimated throughout the IVD. Most critical nutrient concentrations were found within the anterior NP around the mid-transverse plane, with a prediction of an average glucose concentration of 0.8901 mM and pH 6.9349.

The PN-Methodology was embedded in a 3D Agent-based (AB) model (Netlogo (36) version 6.0.2) that simulated a NP environment consisting of 4000 agents (diameter 10µm) and a volume of $1mm^3$. Agents represent NP cells at an average density (37). Agents were randomly placed into the volume and immobile throughout simulations. Each NP cell obtained one out of four cell states; immunonegative or immunopositive for IL1β, for TNF-α or for both IL1β&TNF-α. The setup of the multicellular environment is explained in detail in our previous publication (1). In short: based on the global amount of IL1β and TNF-α mRNA expression provided by the PN-Methodology (ranging between 0 and 1, see Supplementary material 1, PN-Systems II and III), the percentage of immunopositive cells for either IL1β or TNF-α was determined according to the external stimulus concentrations. Corresponding numbers of cells immunopositive for either IL1β or TNF-α were distributed in randomly placed cell clusters. Within regions where IL1β and TNF-α immunopositive cell clusters overlap, cells were considered to be immunopositive for both, IL1β&TNF-α. The minimal calculation time step was 1h.

<u>Results</u>: Early degenerated conditions led to similar CA profiles for cells not exposed to proinflammatory cytokines and cells immumopositive for IL1β as found for non-degenerated conditions but caused a worse catabolic shift under TNF-α presence, under both, gravity and exposure to microgravity. The catabolic shift in cells exposed to TNF-α consists of a downregulation of Col-II and an upregulation of MMP3 and ADAMTS4. In contrast, Agg mRNA expression is similar for non-degenerated and early degenerated conditions, independently from the CS (Figure S7.1 vs. Figure S8.1).

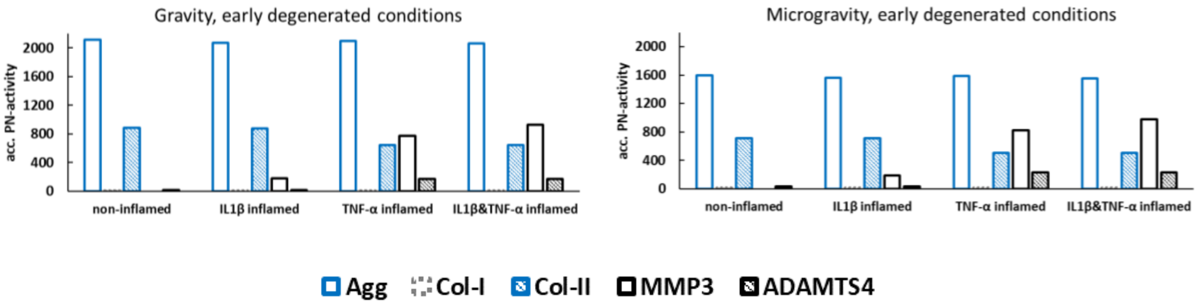


Figure S8.1: estimated CA profiles for an average daily moving habit over six months compared to exposure to microgravity for the same period of time. Simulation of an early degenerated nutrient environment.

The percentage of cells immunopositive for IL1β is maintained around 15-16% for all simulated conditions, whilst TNF-α proinflammatory environments rise from around 10-11% for non-degenerated to 15-16% under early degenerated conditions (Table S8.1).

Table S8.1: predicted, proinflammatory environments within the AB-model for non-degenerated and early degenerated conditions at gravity and under exposure to microgravity. Cells immunopositive for both, IL1β&TNF-α were added to the respective percentages of both, TNF-α and IL1β (see (1)). Percentages of cells immunopositive for TNF-α and IL1β are rounded and based on three model runs for each condition.

| | | IL1β | TNF-α |
|---|---|---|---|
| Non-degenerated | Gravity | ~15.5% | ~10.5% |
| | Microgravity | ~15.5% | ~10.5% |
| Early degenerated | Gravity | ~16% | ~15.5% |
| | Microgravity | ~16% | ~16% |

Discussion: the effect of early degeneration on tissue proteins is small (with the exception of Col-II downregulation under TNF-α presence), which is in agreement with the generally slow development of this disease. However, the change in nutrient concentrations caused an elevated TNF-α presence, leading to a catabolic shift of CA of cells exposed to TNF-α and to an elevated percentage of cells immunopositive for TNF-α (Table S8.1). Hence, catabolic shifts might be locally enhanced within the NP under early degenerated conditions.

Overall, predicted proinflammatory environments lie within plausible ranges, albeit they still lie within ranges as expected in non-degenerated NP, i.e. 17% ± 7%, for IL1β and 16% ± 7%, respectively, for TNF-α (27). Hence, the currently estimated percentage of cells influenced by TNF-α or IL1β might be conservative and, consequently, as well the catabolic shifts of the CA profiles of those cells are conservative estimations.

Current results might suggest TNF-α immunopositivity to play a role under early degenerated conditions. However, the prediction of proinflammatory cytokine expression underlies uncertainties, given that the effect of loading is largely unknown. Moreover, critical factors such as effects between the proinflammatory cytokines (1) or possible endogenous triggers of inflammation such as ECM breakdown products (38) (i.e. damage-associated molecular patterns) were not considered, which coincides with an overall conservative prediction of inflammation.

#### Supplementary material 9

##### Additional analysis of the performance of the PN-Methodology for user-defined conditions around the α-β-threshold

Methods: To investigate the model performance around the α-β-threshold, one loading parameter was set to respective minimal values, i.e. either 0.1 MPa or 0 Hz, whilst the other one was varied, covering the range around the α-β-threshold of the generic functions. Hence, a static condition (0 Hz) was simulated under varying loading conditions of 0.90 MPa, 0.95 MPa, 1.00 MPa, 1.05 MPa and 1.10 MPa and dynamic conditions were simulated under a constant load of 0.10 MPa at 2.90 Hz, 2.95 Hz, 3.00 Hz, 3.05 Hz and 3.10 Hz (note that the minimal step increase of loading parameters was set to 0.05). Thereby, the loading conditions 0.90 MPa, 0.95 MPa and 2.90 Hz, 2.95 Hz, 3.00 Hz, respectively, were initally anabolic, whilst higher loading conditions caused catabolic responses (Figure S9.1). Optimal nutrient conditions (5 mM glucose, pH 7.1) were simulated, and the response of non-inflamed cells was plotted.

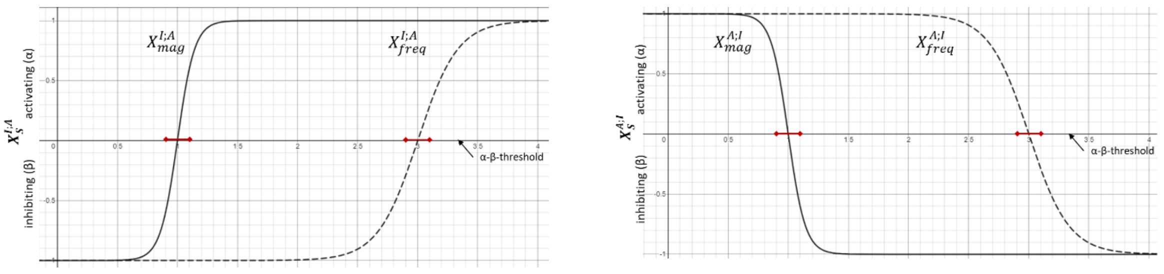


Figure S9.1: generic functions reflecting I;A (left) and A;I (right) evolutions, with critical ranges around the α-β-threshold marked as red bars.

Results: Model predictions were influenced by the steepness of the sigmoidal slope and the impact of the respective stimulus on a specific CA. Hence, the steeper the slope, the shorter the time range of incoherent predictions. (Incoherent predictions refer to incorrect results that might appear due to time-dependent effects within the first hours of calculations. For example, the MMP3 mRNA expressions at 0 Hz and 0.1 MPa; after 2h of persisting loading, the model predicts lowest PN-activity at 1.05 MPa and 3.05 Hz, respectively (Figure S9.2).) Predictions based on the A;I generic function, i.e. tissue protein expressions, quickly converged to the same PN-activity, independently of the initial value of the stimulus (Figure S9.2, Agg, Col-II). In contrast, predictions for I;A generic functions (Col-I and protease expressions) converged much slower (Figure S9.3). Hence, the PN-activity of I;A generic functions initially converge to two different PN-activities over the first hours of simulation (pronounced in Figure S9.2, Col-I, ADAMTS4), depending on their input range (i.e. anabolic or catabolic). Predictions at 1.00 MPa led to systematically erroneous results (Figure S9.2, left column).

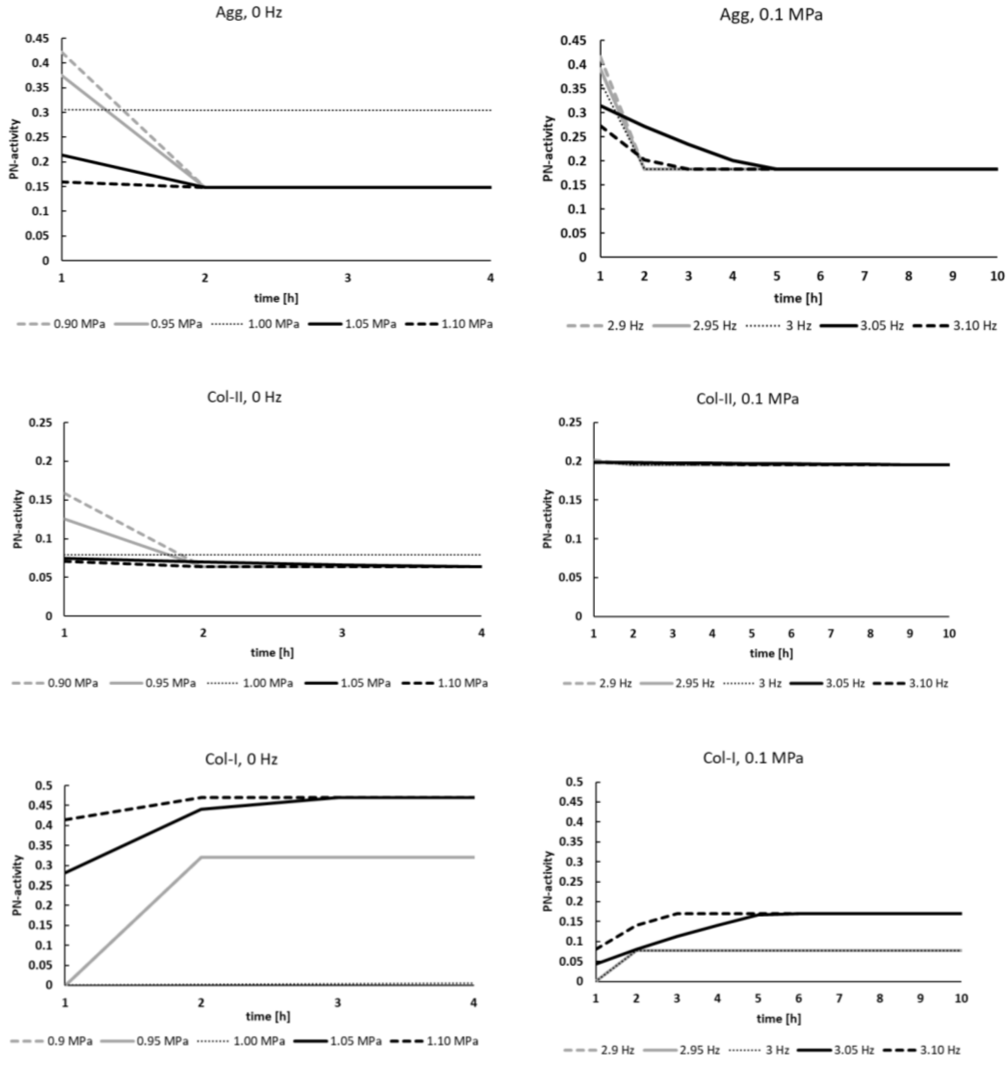

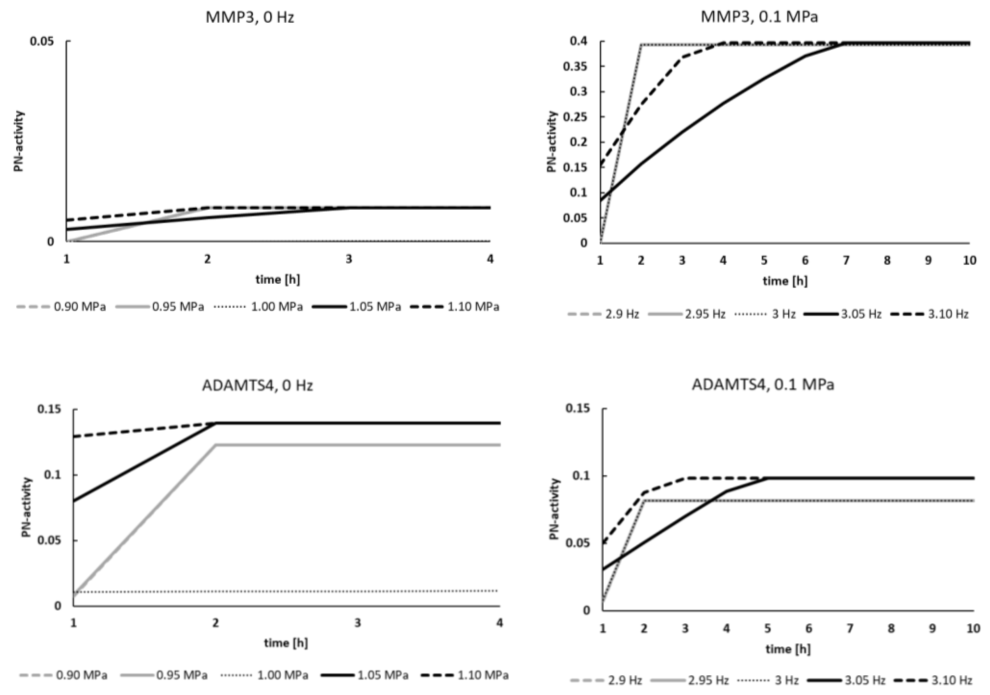


Figure S9.2: predictions around the α-β-threshold of the generic functions of mag (left) and freq (right). Note that predictions at 0.9 and 0.95 MPa are identical for Col-I and MMP3 mRNA expressions and almost identical for ADAMTS4 mRNA expression at 0 Hz (left column). Likewise, predictions at 2.90 Hz, 2.95 Hz and 3.00 Hz have the same values for Col-I and MMP3 mRNA expressions and almost identical values in case of ADAMTS4 mRNA expressions at 0.1 MPa (right column).

Discussion: the influence of the steepness of the sigmoidal shape is not surprising, given that the time sensitivity is based on the generic functions. Hence, a less steep slope includes a smaller effect of time in each time step, which leads to a slower evolution of the CA over time. Stimulus doses closest to the α-β-threshold are 1.00 MPa and 3.00 Hz, respectively (Figure S9.1). Whilst the prediction of 1.00 MPa generally led to incorrect predictions (Figure S9.2, left column), predictions around 3.00 Hz did not show any peculiar behavior (Figure S9.2, right column). The reason is that 3.00 Hz is still considered as an anabolic stimulus by the algorithm, whilst 1.00 MPa is already slightly catabolic. Hence, according to the setup of time sensitivity, anabolic stimuli closely to the α-β-threshold rapidly lose their anabolism, whilst the programming of the latency time assumes that weak catabolic stimulus doses are longer tolerated than strong catabolic stimulus doses. This explains why PN-activities after short time periods might generally lead to incoherent results, e.g. the effect of 3.05 MPa on

Agg mRNA expression (Figure S9.2, “Agg, 0 Hz”) after two hours show a higher PN-activity than any other condition. To overcome such limitations, it is generally recommended to approximate the PN activity by an extrapolation of results calculated at higher and lower stimulus doses.

Whilst the effect of time on CA based on A;I generic functions, i.e. Agg and Col-II mRNA expressions, converge to same PN-activities within a short period of time (with exception of the previously addressed predictions at 1.00 MPa), mRNA expression based on I;A generic functions converge to two different levels of PN-activities, depending on the anabolic or catabolic character of the initial stimulus dose. Thereby, initially anabolic loading led to a generally lower catabolic response than initially catabolic loads (Figure S9.2). This difference is caused by the first fraction of the inhibiting term. The weight that this fraction provides to the inhibiting term is different from the weight within the activating portion of the PN-Equation. This is considered to be a limitation of the PN-Equation. However, qualitatively the results are coherent; initially anabolic values remain more anabolic over time. The effect vanishes as soon as the inhibiting term of the PN-Equation becomes 0, which is subsequently demonstrated by plotting the evolution of Col-I and Agg mRNA expressions over time (Figure S9.3). Note that the freq was elevated from 0 Hz to 2 Hz, to accelerate the decrease of the inhibiting term of the PN-Equation to zero.

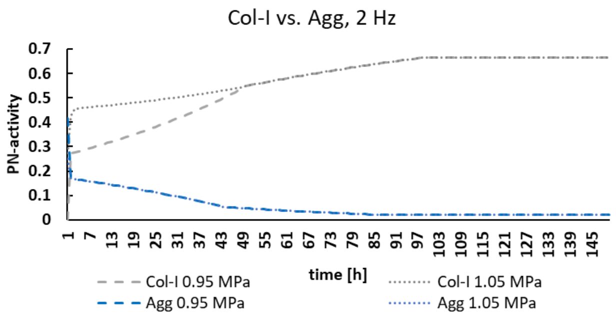


Figure S9.3: evolution of initially anabolic (0.95 MPa) and catabolic (1.05 MPa) mag doses over time on the example of Col-I and Agg. The same PN-activity for Col-I is obtained after 50 h of stimulus exposure. In contrast, the PN-activity of Agg exposed to different mag are already reached after 4 h stimulus exposure.

### 10 References


1. Baumgartner L, Sadowska A, Tio L, González Ballester MA, Wuertz-Kozak K, Noailly J. Evidence-based Network Modelling to Simulate Nucleus Pulposus Multicellular Activity in different Nutritional and pro-Inflammatory Environments. Front Bioeng Biotechnol. 2021;9:1–17.

2. MacLean JJ, Lee CR, Alini M, Iatridis JC. Anabolic and catabolic mRNA levels of the intervertebral disc vary with the magnitude and frequency of in vivo dynamic compression. Journal of orthopaedic Research. 2004;22:1193–200.

3. Gawri R, Rosenzweig DH, Krock E, Ouellet JA, Stone LS, Quinn TM, et al. High mechanical strain of primary intervertebral disc cells promotes secretion of inflammatory factors associated with disc degeneration and pain. Arthritis Res Ther [Internet]. 2014;16(1):R21.

4. Dudli S, Haschtmann D, Ferguson SJ. Fracture of the vertebral endplates, but not equienergetic impact load, promotes disc degeneration in vitro. Journal of Orthopaedic Research. 2012;30(5):809–16.

5. Walter B, Korecki C, Purmessur D, Roughley P, Michalek A, Iatridis J. Complex Loading Affects Intervertebral Disc Mechanics and Biology. Osteoarthritis Cartilage. 2012;19(8):1011–8.

6. Rinkler C, Heuer F, Pedro MT, Mauer UM, Ignatius A, Neidlinger-Wilke C. Influence of low glucose supply on the regulation of gene expression by nucleus pulposus cells and their responsiveness to mechanical loading. Journal of Neurosurgery: Spine J Neurosurg Spine [Internet]. 2010;13:535–42.

7. Gilbert HTJ, Hodson N, Baird P, Richardson SM, Hoyland JA. Acidic pH promotes intervertebral disc degeneration: Acid-sensing ion channel -3 as a potential therapeutic target. Sci Rep [Internet]. 2016;6(1):1–12.

8. Neidlinger-Wilke C, Mietsch A, Rinkler C, Wilke HJ, Ignatius A, Urban J. Interactions of environmental conditions and mechanical loads have influence on matrix turnover by nucleus pulposus cells. Journal of Orthopaedic Research (J Orthop Res). 2012;30(1):112–21.

9. Le Maitre CL, Freemont AJ, Hoyland JA. The role of interleukin-1 in the pathogenesis of human intervertebral disc degeneration. Arthritis Res Ther. 2005;7(4):R732–45.

10. Mendoza L, Xenarios I. A method for the generation of standardized qualitative dynamical systems of regulatory networks. Theoretical biology & medical modelling (Theor Biol Med Model). 2006;3:13.

11. Chan SCW, Ferguson SJ, Gantenbein-Ritter B. The effects of dynamic loading on the intervertebral disc. European Spine Journal. 2011;20:1796–812.

12. MacLean JJ, Lee CR, Alini M, Iatridis JC. The effects of short-term load duration on anabolic and catabolic gene expression in the rat tail intervertebral disc. Journal of Orthopaedic Research. 2005;23(5):1120–7.

13. Belavy DL, Adams M, Brisby H, Cagnie B, Danneels L, Fairbank J, et al. Disc herniations in astronauts: What causes them, and what does it tell us about herniation on earth? European Spine Journal. 2016;25(1):144–54.

14. Purmessur D, Walter B a, Roughley PJ, Laudier DM, Hecht a C, Iatridis J. A role for TNFα in intervertebral disc degeneration: a non-recoverable catabolic shift. Biochem Biophys Res Commun. 2013 Mar;433(1):151–6.

15. NASA. Astronauts answer student questions. 2011. p. National Aeronautics and Space Administration.

16. Wilke HJ, Neef P, Caimi M, Hoogland T, Claes LE. New in vivo measurements of pressures in the intervertebral disc in daily life. Spine (Phila Pa 1976) [Internet]. 1999;24(8):755–62.

17. Pachi A, Ji T. Frequency and velocity of people walking. The Structural Engineer. 2005;36–40.

18. Vernillo G, Giandolini M, Edwards WB, Morin JB, Samozino P, Horvais N, et al. Biomechanics and Physiology of Uphill and Downhill Running. Sports Medicine. 2017;47:615–29.

19. Stokes IAF, Iatridis JC. Mechanical conditions that accelerate intervertebral disc degeneration: Overload versus immobilization. Spine (Phila Pa 1976). 2004;29(23):2724–32.

20. Luoma K, Riihimäki H, Luukkonen R, Raininko R, Viikari-Juntura E, Lamminen A. Low Back Pain in Relation to Lumbar Disc Degeneration. Spine (Phila Pa 1976). 2000;25(4):487–92.

21. Takatalo J, Karppinen J, Näyhä S, Taimela S, Niinimäki J, Blanco Sequeiros R, et al. Association between adolescent sport activities and lumbar disk degeneration among young adults. Scand J Med Sci Sports. 2017;27:1993–2001.

22. Belavý DL, Quittner MJ, Ridgers N, Ling Y, Connell D, Rantalainen T. Running exercise strengthens the intervertebral disc. Sci Rep. 2017;7:1–8.

23. Videman T, Sarna S, Battié MC, Koskinen S, Gill K, Paananen H, et al. The Long-Term Effects of Physical Loading and Exercise Lifestyles on Back-Related Symptoms, Disability, and Spinal Pathology Among Men. Spine (Phila Pa 1976). 1995 Mar;20(6):699–709.

24. Nachemson A, Elfström G. Intravital dynamic pressure measurements in lumbar discs. Vol. 2, Scandinavian Journal of Rehabilitation and Medication. 1970. p. 1–40.

25. Hangai M, Kaneoka K, Hinotsu S, Shimizu K, Okubo Y, Miyakawa S, et al. Lumbar intervertebral disk degeneration in athletes. American Journal of Sports Medicine. 2009;37(1):149–55.

26. Mitchell UH, Bowden JA, Larson RE, Belavy DL, Owen PJ. Long-term running in middle-aged men and intervertebral disc health , a cross-sectional pilot study. PLoS One. 2020;15(2):e0229457.

27. Molinos M, Almeida CR, Caldeira J, Cunha C, Goncalves RM, Barbosa MA. Inflammation in intervertebral disc degeneration and regeneration. J R Soc Interface. 2015;12(104):20141191.

28. Sivan SS, Tsitron E, Wachtel E, Roughley PJ, Sakkee N, Van Der Ham F, et al. Aggrecan turnover in human intervertebral disc as determined by the racemization of aspartic acid. Journal of Biological Chemistry. 2006;281(19):13009–14.

29. Sivan SS, Wachtel E, Tsitron E, Sakkee N, Van Der Ham F, DeGroot J, et al. Collagen turnover in normal and degenerate human intervertebral discs as determined by the racemization of aspartic acid. Journal of Biological Chemistry. 2008;283(14):8796–801.

30. Baumgartner L, Wuertz-Kozak K, Le Maitre CL, Wignall F, Richardson SM, Hoyland J, et al. Multiscale regulation of the intervertebral disc: Achievements in experimental, in silico, and regenerative research. Int J Mol Sci. 2021;22(2):1–42.

31. Le Maitre CL, Hoyland JA, Freemont AJ. Catabolic cytokine expression in degenerate and herniated human intervertebral discs: IL-1beta and TNFalpha expression profile. Arthritis Res Ther [Internet]. 2007;9(4):R77.

32. Garcia KM, Harrison MF, Sargsyan AE, Ebert D, Dulchavsky SA. Real-time Ultrasound Assessment of Astronaut Spinal Anatomy and Disorders on the International Space Station. Journal of Ultrasound in Medicine. 2018;37:987–99.

33. Koy T, Zange J, Rittweger J, Pohle-Fröhlich R, Hackenbroch M, Eysel P, et al. Assessment of lumbar intervertebral disc glycosaminoglycan content by gadolinium-enhanced MRI before and after 21-days of head-down-tilt bedrest. PLoS One. 2014;9(11):e112104.

34. Bailey JF, Miller SL, Khieu K, O´Neill CW, Healey RM, Coughlin DG, et al. From the international space station to the clinic: how prolonged unloading may disrupt lumbar spine stability. Spine J. 2018;18(1):7–14.

35. Ruiz Wills C, Foata B, González Ballester MÁ, Karppinen J, Noailly J. Theoretical Explorations Generate New Hypotheses About the Role of the Cartilage Endplate in Early Intervertebral Disk Degeneration. Front Physiol [Internet]. 2018;9:1–12. Available from: https://www.frontiersin.org/article/10.3389/fphys.2018.01210/full

36. Wilensky U. NetLogo. 1999. p. NetLogo. http://ccl.northwestern.edu/netlogo/.

37. Maroudas A, Stockwell RA, Nachemson A, Urban J. Factors involved in the nutrition of the human lumbar intervertebral disc: cellularity and diffusion of glucose in vitro. Journal of Anatomy (J Anat) [Internet]. 1975;120(Pt 1):113–30.

38. Medzhitov R. Origin and physiological roles of inflammation. Nature. 2008;454:428–35.
